# Cohabiting microbes can impact one another’s mutation rates creating a mutational eco-evo feedback loop

**DOI:** 10.64898/2026.11.06.764923

**Authors:** R Green, E Wright, R Palmer, R Tank, E Tonge, J Scaife, A Marquiegui-Alvaro, C G Knight, R Krašovec

**Affiliations:** School of Natural Sciences, Faculty of Science & Engineering, University of Manchester; School of Biological Sciences, Faculty of Biology, Medicine & Health, University of Manchester; Department of Physics and Astronomy, Faculty of Science & Engineering, University of Manchester; Department of Zoology, Faculty of Science, University of British Columbia; Department of Chemistry, and Manchester Institute of Biotechnology (MIB), University of Manchester

**Keywords:** Mutation rate, eco-evo feedback, mutation rate plasticity, co-culture, E. coli, E. faecalis, P. aeruginosa

## Abstract

Eco-evolutionary feedbacks are typically thought to require an ecological change driving Darwinian selection. However, ecology can drive other evolutionary mechanisms too. In many microbial species mutation rates are elevated at low population density, mediated by nutrient availability and reactive oxygen species (ROS). These findings from monoculture populations led us to hypothesise that the presence and density of a cohabiting species will also affect mutation rates. Using Escherichia coli as a focal species we explore the mutational effects on both parties of co-culture with either Enterococcus faecalis or Pseudomonas aeruginosa. These species co-occur in multiple clinical settings and have different abilities to degrade ROS. In both co-culture systems we find evidence that the presence of a stronger ROS degrader reduces mutation rates in the weaker ROS degrading species. At the same time increasing population density of the weaker ROS degrader increases mutation rates in the stronger ROS degrading species. Simultaneous mutation rate estimation in co-cultured species raises multiple experimental challenges addressed here. Despite these challenges, we find evidence for a novel form of eco-evo feedback in which the environment shapes the supply of mutational variation to the population. This in turn feeds back to alter the chance of evolutionary rescue for one species or another during a subsequent environmental change. Such effects potentially matter not only for our fundamental understanding of the interplay between ecology and evolution, but for the treatment of polymicrobial infections in which rates of resistance evolution may be altered by the cross-species mutational effects of cohabiting species.

## Introduction

Where evolutionary and ecological dynamics occur on similar timescales, notably in microbial communities, eco-evolutionary feedbacks may be observed. Eco-evolutionary feedbacks require that “populations alter their environment…and that those changes in the environment feed back to influence…subsequent evolution” [1]. Any effect on subsequent evolution is generally expected to act via changes in selection pressures [2]. However, natural selection is not the only evolutionary mechanism affected by ecological factors, mutation is another evolutionary force known to be impacted by the environment. Mutation rates vary due to both genetic and environmental factors. Within a single genotype mutation rates have been shown to vary with environmental factors such as stress [3], nutrient availability [4, 5], oxygen [5], temperature [6, 7] and population density [8]. Variation in mutation rate shapes evolutionary trajectories [9], for example, elevated mutation rates, under strong selective pressures such as antibiotic treatment, accelerate the emergence and spread of antimicrobial resistance (AMR) [10–12]. Environmentally plastic mutation rates have the potential to aid adaptation [13, 14], although it is unclear how environmentally induced variation plays out in natural populations. Most studies of mutation rate variation focus on single species systems, despite microbes commonly inhabiting diverse polymicrobial communities [15]. In multispecies environments, co-resident species may mediate effects on mutation rate, for example by excreting or degrading stressor or mutagenic molecules in the shared environment. Understanding these interspecies feedbacks via mutation rates is therefore essential for predicting eco-evolutionary dynamics beyond laboratory monocultures.

Mutation rate plasticity in mixed species communities is not well studied. While it is known that interspecies interactions can influence evolutionary genetic trajectories [16–18], these studies do not explicitly separate the effects of the community on mutation from its effects on selection. Cross-species mutagenic effects have however been explicitly considered in the case of cytosine deaminase toxins used in bacterial warfare. Secretion of these mutagenic toxins by *Burkholderia cenocepacia* has been demonstrated to increase mutation rates in surviving *Escherichia coli* competitors [19]. Further suggestive evidence for cross-species mutagenic effects has been found in multiple bacterial species however, these previous studies fail to accurately estimate mutation rates [20] or to account for the impact of multicellular aggregation on these estimates [21].

Despite limited work on cross-species mutational effects, within-species community effects on mutagenesis have been well characterised in the case of Density Associated Mutation rate Plasticity (DAMP). DAMP is a widespread trait across domains of life, characterised by populations at higher cell density experiencing reduced per-genome per-generation mutation rates (a separate mechanism from stress-induced mutagenesis) [8, 22]. In this context cross-strain mutation rate effects have been characterised in multiple *E. coli* strain pairs [23, 24]. Collective hydrogen peroxide degradation provides a mechanistic basis for DAMP in single species cultures of *E. coli* [24]. In contrast, *Pseudomonas aeruginosa,* despite being closely related, does not show a substantial DAMP response [8]. This difference is likely due to *P. aerguinosa*’s high hydrogen peroxide degradation capacity and low outer membrane permeability, which provide resistance to oxidative damage even at low cell densities [24–26]. Therefore, the environmental responsiveness of mutation rates may evolve as a result of selection on related traits, such as membrane permeability and protection from reactive oxygen species (ROS).

In addition to being implicated in the DAMP phenotype ROS, including hydrogen peroxide, are common stressors that vary in production and degradation across bacterial species [25]. The down-regulation of ROS degradation enzymes responsible for degrading hydrogen peroxide has previously been implicated in the increase of *P. aeruginosa* mutation rates during biofilm-co-culture with the fungus *C. albicans* [20]. ROS also mediate cross-protection and competition within microbial communities [27–29]. We therefore hypothesise that ROS will mediate interspecies as well as intraspecies mutation rate plasticity depending on the hydrogen peroxide production and detoxification capacities of the species involved. We further hypothesise that this effect will impact the relative chances of evolutionary rescue [30] in cohabiting microbes.

In this study we address mutation rates in mixed-species environments by exploring cross-species mutational effects in two co-culture systems: *P. aeruginosa* + *E. coli* and *E. coli* + *E. faecalis*. Pairings were selected for three key reasons: 1) These species display highly contrasting ROS degradation and production phenotypes, which are known determinants of mutation rate plasticity [24, 25, 31]; 2) Mutation rate plasticity in *E. coli* is relatively well characterised, providing a strong framework for investigating novel aspects of mutation rate plasticity [8, 22–24, 32]; and 3) All three species rank within the top 12 pathogens responsible for AMR-associated mortality in 2019 [33] and are frequently co-isolated in polymicrobial infections [34–42]. The common co-occurrence of the species studied here in clinical infections provides a strong motivation to understand how their interactions influence mutational processes, particularly the likelihood of spontaneous antimicrobial resistance (AMR).

In contrast to *P. aeruginosa*, *E. faecalis* produces superoxide at rates >150-fold higher than *E. coli* [31, 43] and is less efficient at detoxifying hydrogen peroxide than either *E. coli* or *P. aeruginosa* [25]. These three species form a gradient of ROS detoxification capacity from low in *E. faecalis* to high in *P. aeruginosa.* We predict that species that more efficiently remove environmental hydrogen peroxide will reduce the mutation rate of less efficient detoxifiers in co-culture [25]. We therefore estimate mutation rates, using fluctuation assays to rifampicin resistance [44], in mono-and co-culture for *P. aeruginosa* + *E. coli* and *E. coli* + *E. faecalis* pairings, across a range of nutrient-determined population densities.

Using these two co-culture systems, we demonstrate that both the presence and population density of a cohabiting species can alter microbial mutation rates. Through evaluating the validity of our own results, we also highlight the assumptions and analytical considerations necessary for quantifying mutation rates in mixed-species environments. Through its effects on evolutionary rescue (the chances of one species or the other surviving the sudden imposition of lethal selection in the fluctuation test used to estimate mutation rates), this study demonstrates a novel form of eco-evo feedback: ecological conditions alter mutation rates leading to differences in standing genetic variation which alter the relative probabilities of survival following an environmental change. This mutation based phenomena differs from currently understood eco-evo feedbacks which are generally based on changes in selective pressures rather than mutational processes. This work also represents an important step toward measuring mutation rates in settings more complex than single species monocultures. We provide a foundation for future studies aiming to map how natural variation in mutation rates is driven by interspecies effects and the long term eco-evolutionary consequences of these mutational effects.

## Methods & Materials

### Strains

We used *Escherichia coli* strain K12 MG1655 *attB::Km-gfp* (MV1463) [45], *Enterococcus faecalis* strain ATCC 19433 and *Pseudomonas aeruginosa* strain PA01. All species were sequenced to ≥30x coverage by MicrobesNG with short read illumina sequencing to confirm strain identity. For MV1463 the reference sequences used were U00096.3 [46], L29345.1 [GFP from 47] and the *E. coli* kanamycin resistance gene. For ATCC19433 the reference sequence used was assembly_bb21354536ae4198_1 sourced from https://genomes.atcc.org/genomes/5ea20e0fe37644ed [accessed April 2024 from 48]. For PA01 the reference sequence used was NC_002516.2 [49]. Variants between strains and reference sequences can be found in supplementary data file 11.

### Media

Autoclaved Milli-Q water was used for all media preparations and sterile 0.85% saline (8.5 g/L NaCl) was used for all cell dilutions. LB medium, Davis Minimal (DM) medium, M9 minimal medium, Tetrazolium arabinose/lactose agar medium (TA/TL agar) and Brain Heart Infusion (BHI) recipes are given in supplementary data file 12. In the *E. coli* + *E. faecalis* system for mono and co-culture experiments *E. coli* was plated on TA agar (± rifampicin) while *E. faecalis* was plated on BHI agar supplemented with nalidixic acid to which *E. faecalis* is intrinsically resistant unlike *E. coli* (± rifampicin). In the *P. aeruginosa* + *E. coli* system all cultures were plated on TL agar (± rifampicin) on which *P. aeruginosa* forms large deep red colonies while *E. coli* forms smaller white colonies with a red centre. DM media was used in the *E. coli* + *E. faecalis* system for consistency with previous studies [8, 22–24] while M9 media was used for the *P. aeruginosa* + *E. coli* system as *P. aeruginosa* can metabolise the citrate in DM media, leading to excessively dense cultures. DM media was supplemented with both glucose and casamino acids (CAA), as CAA are required for sufficient *E. faecalis* growth, while M9 was supplemented only with glucose.

### Fluctuation assays

Fluctuation assays were carried out as in [44] with some modifications to allow for co-estimation of mutation rates for both species in co-culture as shown in Figure 1. For all fluctuation assays each species was separately inoculated from glycerol stocks into rich media until exponential phase was reached. This culture was then diluted into minimal media for overnight acclimatisation. For all monoculture and co-culture assays relating to the *Pa* + *Ec* system overnight growth was carried out in M9 media supplemented with the appropriate glucose concentration of the fluctuation assay environment to allow for acclimatisation. Following overnight growth all cultures were standardised to ∼3000 CFU/mL in minimal media. Minimal media was supplemented with variable concentrations of glucose (and casamino acids in the *E. coli* + *E. faecalis* system) to allow the estimation of mutation rates across a variety of population densities. For each fluctuation assay a single large culture (usually 22mL) was divided into 3 N_t_ wells and 13-18 selective wells, these parallel cultures of 1mL (or 1.5mL in the case of some PA01 monoculture assays, allowing for mutation rate estimation at lower population densities) were grown in a deep 96-well plate. For co-culture assays both species were inoculated separately into the same set of wells at a ratio of 990:10µL (*Ef*:*Ec*) or 960:40µL (*Ec:Pa*). For *P. aeruginosa* + *E. coli* the same selective wells were used for both species while for *E. coli* + *E. faecalis* separate selective wells were plated on agar selective for each species. After 24 hours of growth N_t_ wells were appropriately diluted prior to plating on non-selective media while selective wells were plated in their entirety on rifampicin selective agar.

**Figure 1:**
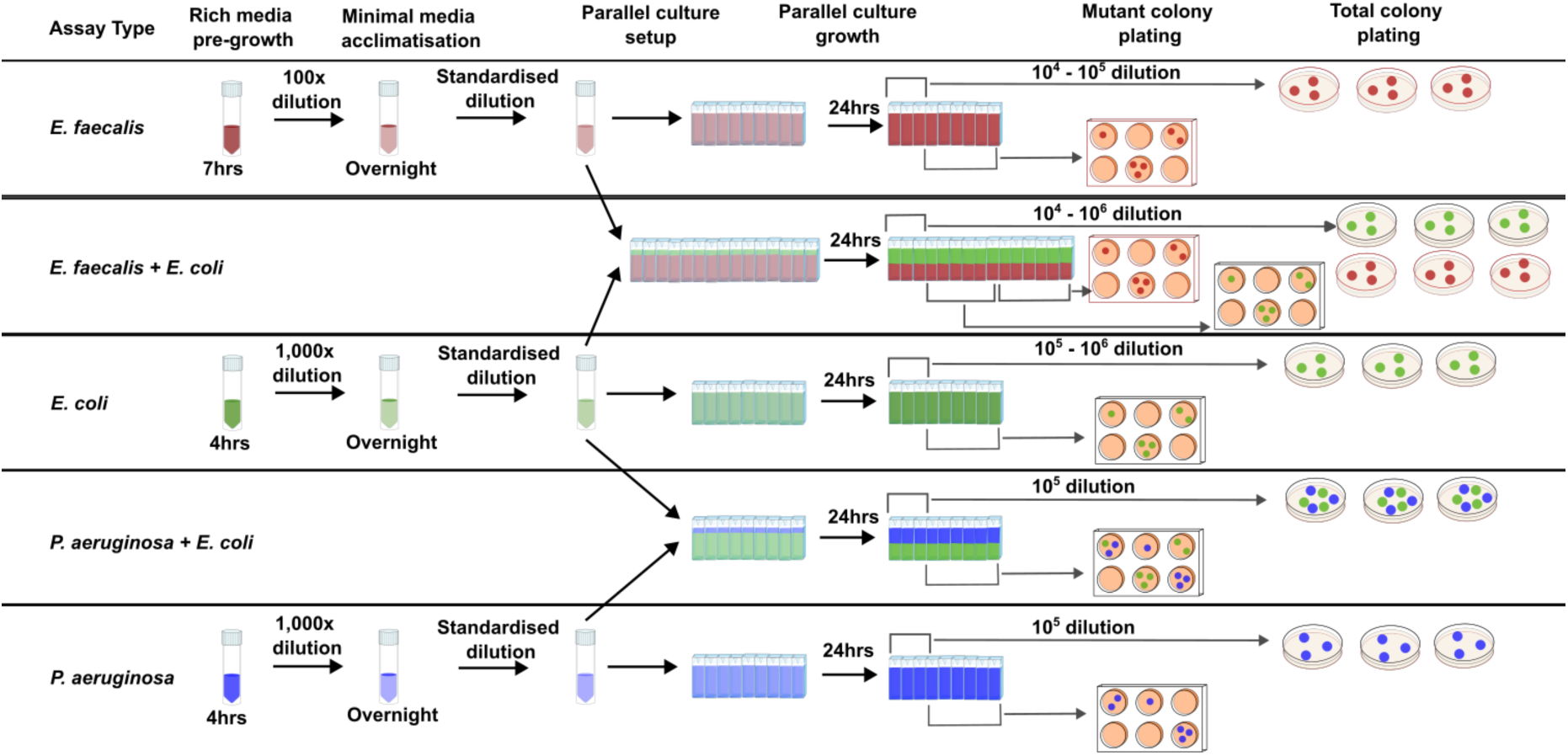
Method overview for mono and co-culture fluctuation assays. Each row shows a fluctuation assay type with headers in the top row indicating experimental phases. Although not shown, *E. coli* monocultures were separately carried out in M9 media for comparison with *P. aeruginosa* experiments and in DM media for comparison with *E. faecalis* experiments. Standardised dilutions were calibrated for each species to a final density of ∼3000 CFU/mL. On the right hand, 6-well plates are rifampicin selective agar while individual plates are non-selective or, in the case of *E. faecalis* plating, contain nalidixic acid to which this species is intrinsically resistant. Vessels shown from left to right represent 50mL falcon tubes, individual wells of a 2mL well-volume deep 96-well plate, 6-well agar plates and 9cm individual agar plates.

### Estimation of mutation rates

The average number of mutational events per culture in each fluctuation assay (*m*) was estimated for each species from the observed rifR mutant CFU counts using the Luria-Delbrϋck maximum likelihood (ML) method implemented using R package ‘flan’ ([50] V1.0). The default expectation of exponential lifetimes (“LD” model) was selected and the maximum likelihood method applied. The occurrence of ‘jackpot’ events were accounted for with a Winsorization parameter of 1024. Mutation rates to rifampicin resistance per generation were calculated by dividing *m* by the final population size ‘N_t_’, as determined by CFU counts on non-selective agar. Where the GF method of co-estimating the fitness effect of rifampicin resistance (*ρ*) along with *m* failed then *ρ* was estimated using Statistical Model 6 and the mutation rate estimation was re-run with this estimate of *ρ* supplied in the ‘fitness’ argument.

### Flow cytometry

#### Sample preparation

*E. coli* and *P. aeruginosa* monocultures and co-cultures were grown under fluctuation assay conditions in M9 media supplemented with either 250mg/L or 1000 mg/L glucose. For each species, mono vs co-culture & glucose combination, nine biological replicates were prepared Table 1. These were divided into three treatment groups: heat-treated, methanol-treated (both serving as positive controls for cell death) and untreated, with three biological replicates per group. For the heat-treated group, samples were incubated on a 65°C heat block for 30 minutes. Methanol-treated samples were first centrifuged at 8000 rpm for 5 minutes to pellet the cells. The supernatant was discarded, and 1mL of 100% methanol was added. Samples were vortexed briefly and incubated at 4°C for 15 minutes. After treatment, all samples were centrifuged at 8000 rpm for 5 minutes, the supernatant was removed, and the pellets were resuspended in 1mL of phosphate-buffered saline (PBS). Propidium iodide (PI) was added to all 1mL samples at a final concentration of 1µg/mL. Samples were incubated at room temperature for 15 minutes in the dark. All samples were then centrifuged again at 8000 rpm for 5 minutes, the supernatant was removed and 300µL of PBS was added to resuspend the cells. Finally, 250µL of each sample was transferred into wells of a U-bottomed 96-well plate for flow cytometry analysis.

#### Sample measurement

Samples were analysed using a CytExpert flow cytometer. A blank PBS well was first run to adjust forward and side scatter (FSC and SSC) gating, excluding background noise and debris to ensure only bacteria-sized particles were included in the analysis. The phycoerythrin (PE) channel was used to detect PI fluorescence, with a default gain of 300. The fluorescein isothiocyanate (FITC) channel was used to detect *E. coli* green fluorescent protein (GFP) fluorescence, with a default gain of 1500. Wells were run at 17µL/min, and stop conditions were set to 2 minutes or until 18,000 bacteria-positive events were recorded. Data was exported and analysed using the flowCore R package, v2.14.2 [51].

#### Microscopy

Monocultures and co-cultures were grown under fluctuation assay conditions in 1mL parallel cultures. 5µL of each sample was placed on a glass slide with a coverslip. Cells were imaged with a 100X oil objective using phase contrast and GFP fluorescence channels to differentiate *E. coli* (GFP-tagged) from non-fluorescent *P. aeruginosa* in co-culture. Images were exported and analysed using ImageJ [52].

## Data Analysis

All data were analysed in R. Packages nlme [53], MASS [54], emmeans [55] and car [56] were used for linear mixed-effects models. P values within each fitted model were adjusted using a false discovery rate correction. T-tests related to Figure 5 were carried out as paired 2-tailed tests. See Supplementary Statistics for further details.

The tidyverse [57] packages magrittr [58], plyr [59] and dplyr [60] were used for data handling. R packages ggpubr [61], ggplot2 [62], gridExtra [63], ggbeeswarm [64], RColorBrewer [65], scales [66], cowplot [67] and ggblend [68] were used for visualisation.

## Results

### Development of co-culture systems for mutation rate measurement

To assess mutation rates in clinically relevant species pairs with diverse ROS production and degradation phenotypes we developed co-culture systems for *P. aeruginosa* + *E. coli* and for *E. coli* + *E. faecalis* (Figure 2). Both co-culture systems used minimal media with differences based on metabolic requirements of the species pairing (see Methods). The number of generations completed by one species relative to the other species during a 24-hour batch culture (relative fitness, *W*) was higher for the stronger ROS degrader: *P. aeruginosa* has a fitness of 1.32 (95% CI = 1.11 - 1.57) relative to *E. coli* while *E. faecalis* has a fitness of 0.58 (95% CI = 0.388 - 0.872) relative to *E. coli* (Figure S1). We measured mutation rates to rifampicin resistance in these systems using the fluctuation assay. This classic method makes various assumptions about colony formation and mortality [69, 70] whose effects on our results we consider in detail in Supplementary Text 1.

**Figure 2:**
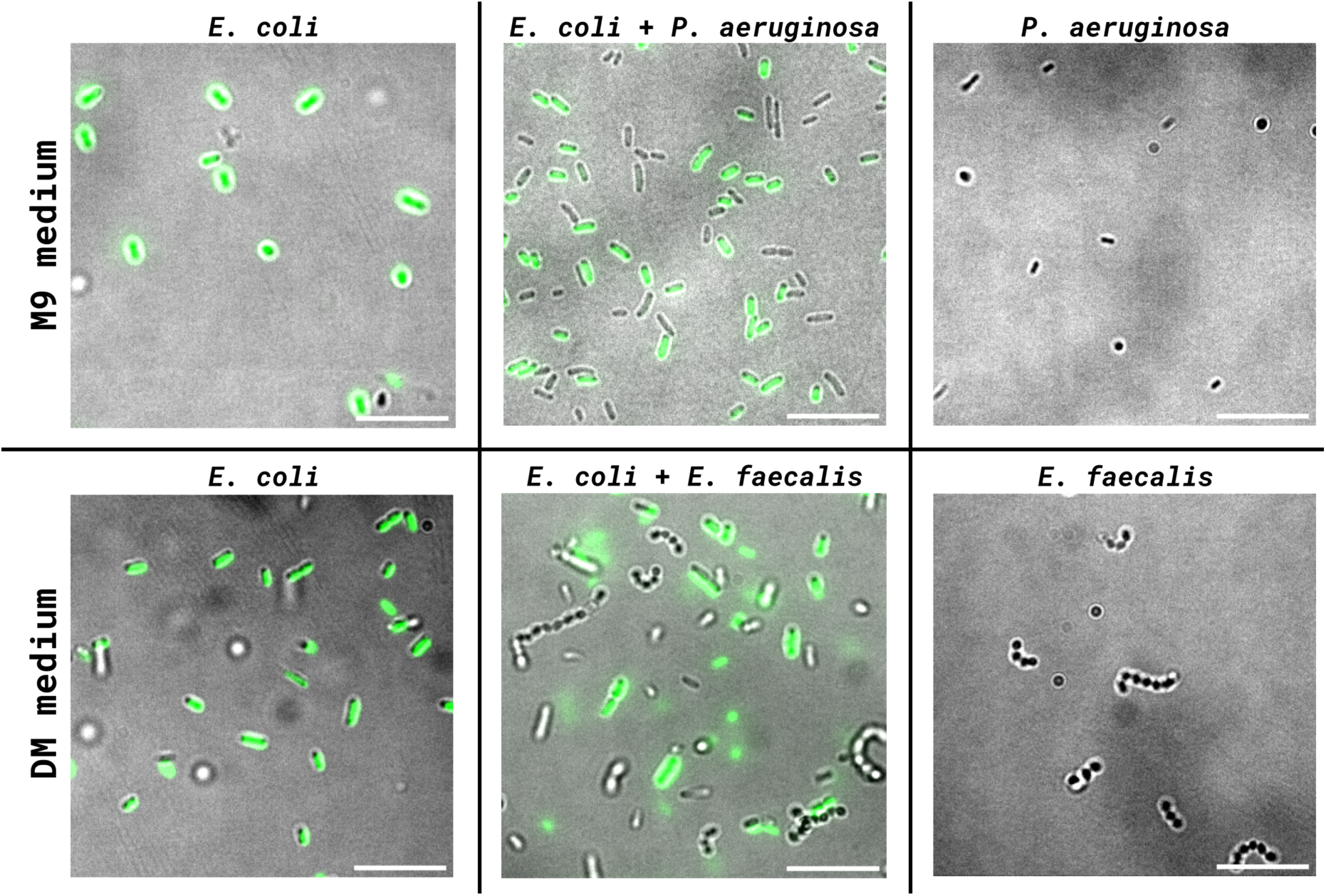
Merged phase contrast and GFP images of *E. coli* and *P. aeruginosa* in monoculture and co-culture (top row) and *E. coli* and *E. faecalis* in monoculture and co-culture (bottom row). The *E. coli* strain used carries the constitutively expressed chromosomal fusion attB::Km-gfp. All cultures were grown as for a fluctuation assay and imaged after 24 hours of growth, this is the time at which cultures are plated to quantify colony forming units (CFUs) in the fluctuation assay (see Methods). Cultures in the top row were grown in M9 medium supplemented with 1,000, 1,000 & 80 mg per L glucose respectively and cultures in the bottom row were grown in DM medium supplemented with 0.15% casamino acids and 500 mg per L glucose. All scale bars are 10µm in length.

### Mutation rates are reduced in the presence of a stronger ROS degrading species

ROS degradation is critical in lowering mutation rates, particularly at high population densities [24, 28]. We therefore hypothesise that species with greater ROS degradation capacity will reduce the mutation rate of a co-cultured species. To test this hypothesis we use three species known for having different ROS degradation capacities: *P. aeruginosa* has higher ROS degradation capacity than *E. coli* and *E. coli* higher than *E. faecalis* [25]. We find that the presence of stronger ROS degraders (*P. aeruginosa* & *E. coli* respectively) significantly reduce mutation rates in the weaker ROS degraders during co-culture in both systems: *E. coli* mutation rates are reduced by 1.8-fold in the presence of *P. aeruginosa* (95% CI: 1.3-2.7, *t* ratio_124_ = 3.16, *P* = 0.0049, Statistical Model 1) and *E. faecalis* mutation rates are reduced by 1.6-fold in the presence of *E. coli* (95% CI: 1.3 - 2, *t* ratio_53_ =-4.2, *P* = 1.20 × 10^−4^, Statistical Model 2). Thus the presence of a cohabiting species can alter microbial mutation rates. The direction of this effect is consistent with an antimutagenic reduction in extracellular ROS concentrations achieved by a stronger ROS degrader.

### Higher population densities of a weaker ROS degrading species are associated with increased mutation rates in a stronger ROS degrading species

In Figure 3 we assessed whether the presence of a co-cultured species is associated with a change in the average mutation rate of a focal species, considering the co-culture treatment as a binary variable. However, each co-cultured population represents a unique environment defined by the specific population densities of both species (Figure 4A&C). We therefore ask if mutation rates depend on those specific population densities, expecting to see an increase in mutation rate with increasing densities of lower capability ROS degraders and vice versa. Among *P. aeruginosa* + *E. coli* co-culture experiments we find that *P. aeruginosa* mutation rates are increased 1.4-fold with each doubling of the *E. coli* population density (95% CI: 1.1-1.8, *t* ratio_14_ = 2.9, *P* = 0.04, Figure 4B, Statistical Model 3). *E. coli* mutation rates are however not substantially altered as the density of *P. aeruginosa* increases (0.93-fold change with *Pa* density doubling, 95% CI: 0.74 - 1.2, *t* ratio_14_ =-0.4, *P* = 0.69, Statistical Model 3). This result combines with those from Figure 3 to suggest that while *E. coli* mutation rates are uniformly decreased by the presence of *P. aeruginosa*, *P. aeruginosa* mutation rates are increased only at high densities of *E. coli*.

**Figure 3:**
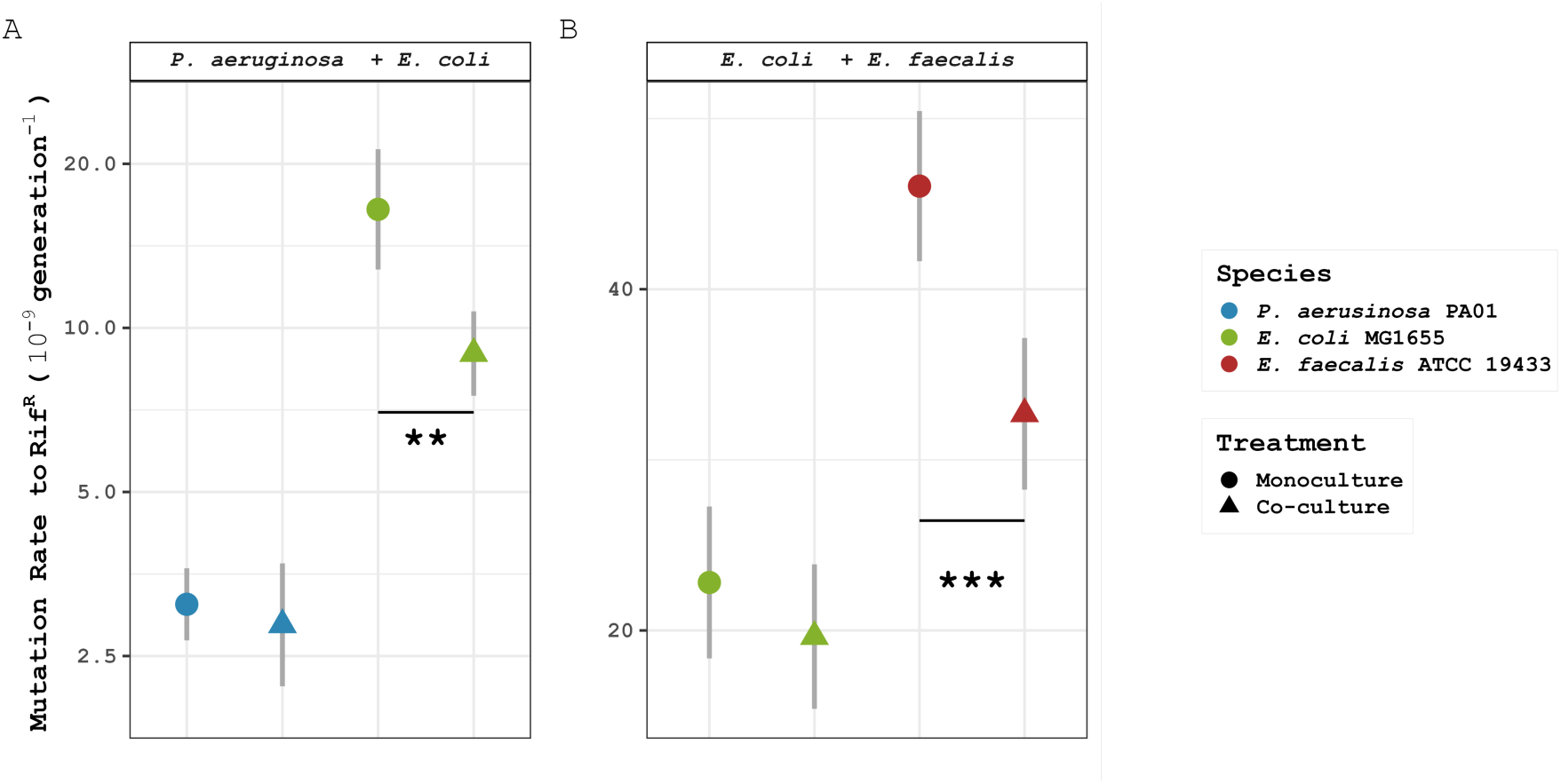
Mutation rate to rifampicin resistance differs between mono and co-culture conditions. **A)** Mutation rates of *P. aeruginosa* (N_mono_ = 90, N_co_ = 29) and *E. coli* (N_mono_ = 9, N_co_ = 31), all experiments in M9 medium with glucose. Mutation rates are estimated at the mean population density of 2.24 × 10^8^ for both species. Raw mutation rate estimates are shown in Figure S2. **B)** Mutation rates of *E. coli* (N_mono_ = 12, N_co_ = 24) and *E. faecalis* (N_mono_ = 12, N_co_ = 24), all experiments in DM medium with glucose and casamino acids. Mutation rates estimated at mean population density by species (*E. coli* = 5.84 × 10^8^, *E. faecalis* = 2.43 × 10^7^). Raw mutation rate estimates are shown in Figure S3. Across panels A&B species are ordered from left to right from strongest to weakest ROS degrading phenotype as measured by [25]. Error bars show ±2 standard errors of the mean. Note log scale y-axes with differing ranges. Despite being measured in different media used for *P. aeruginosa* and *E. faecalis*, the two *E. coli* monoculture estimates are very similar indicating similar mutation rates in these 2 environments.

**Figure 4:**
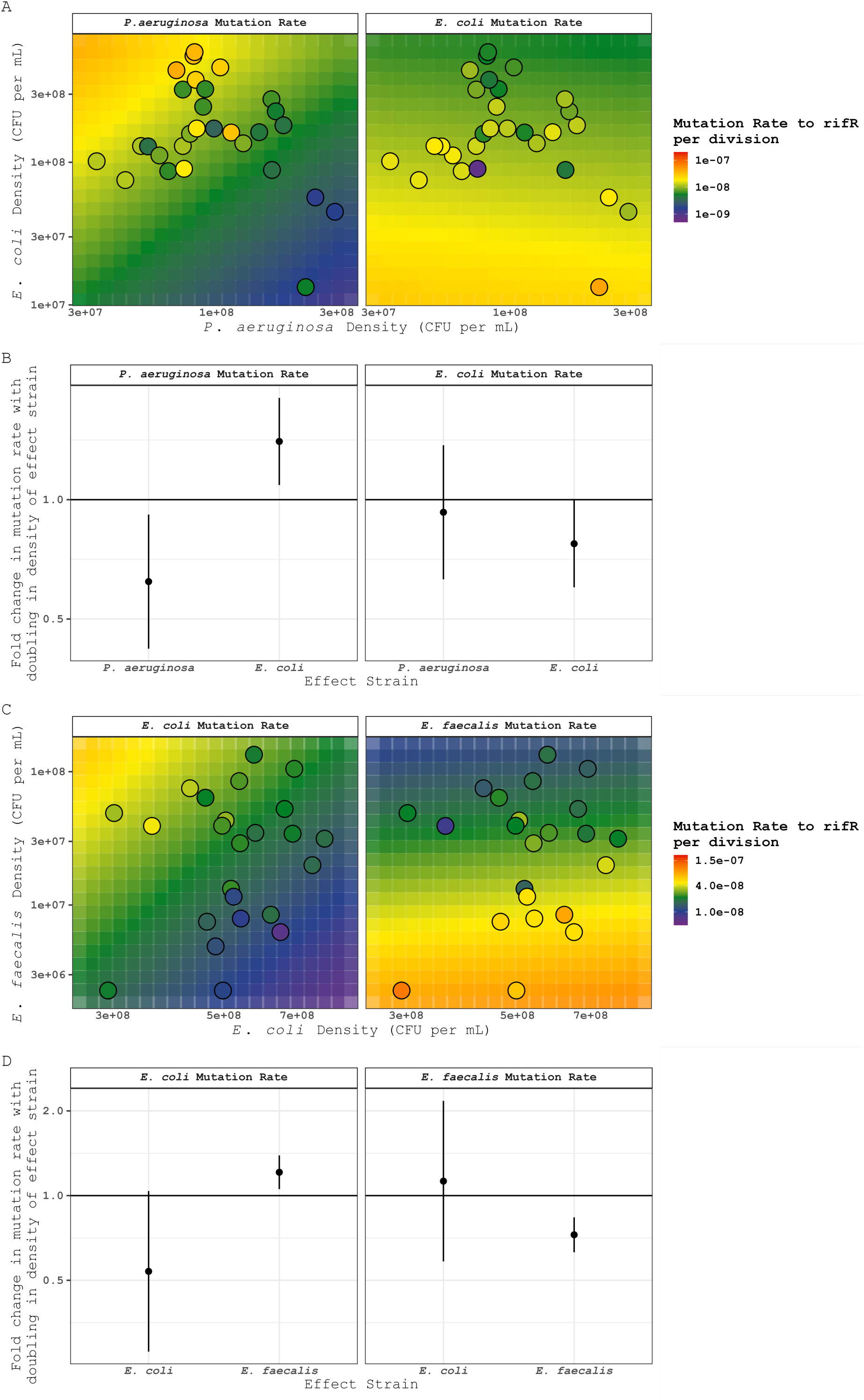
Mutation rates in co-culture as a function of the final population density of both species. A&C) Each panel shows data (circles) and model fitted predictions (background squares) for mutation rates of *P. aeruginosa* and *E. coli* (panel A, left and right respectively) and *E. coli* and *E. faecalis* (panel B, left and right respectively). Note log-log axes with different ranges on x vs y axes for A and C. B&D) The modelled mutational effect of the density of the ‘effect strain’ (shown in the x axis) is plotted for each species (shown in the facet header). Y-axis shows the fold change in mutation rate of the focal species (facet header) with each doubling of the density of the effect strain (x axis). Error bars show 95% confidence intervals. Details of fitted models can be found in Statistical Models 3 & 4.

**Figure 5:**
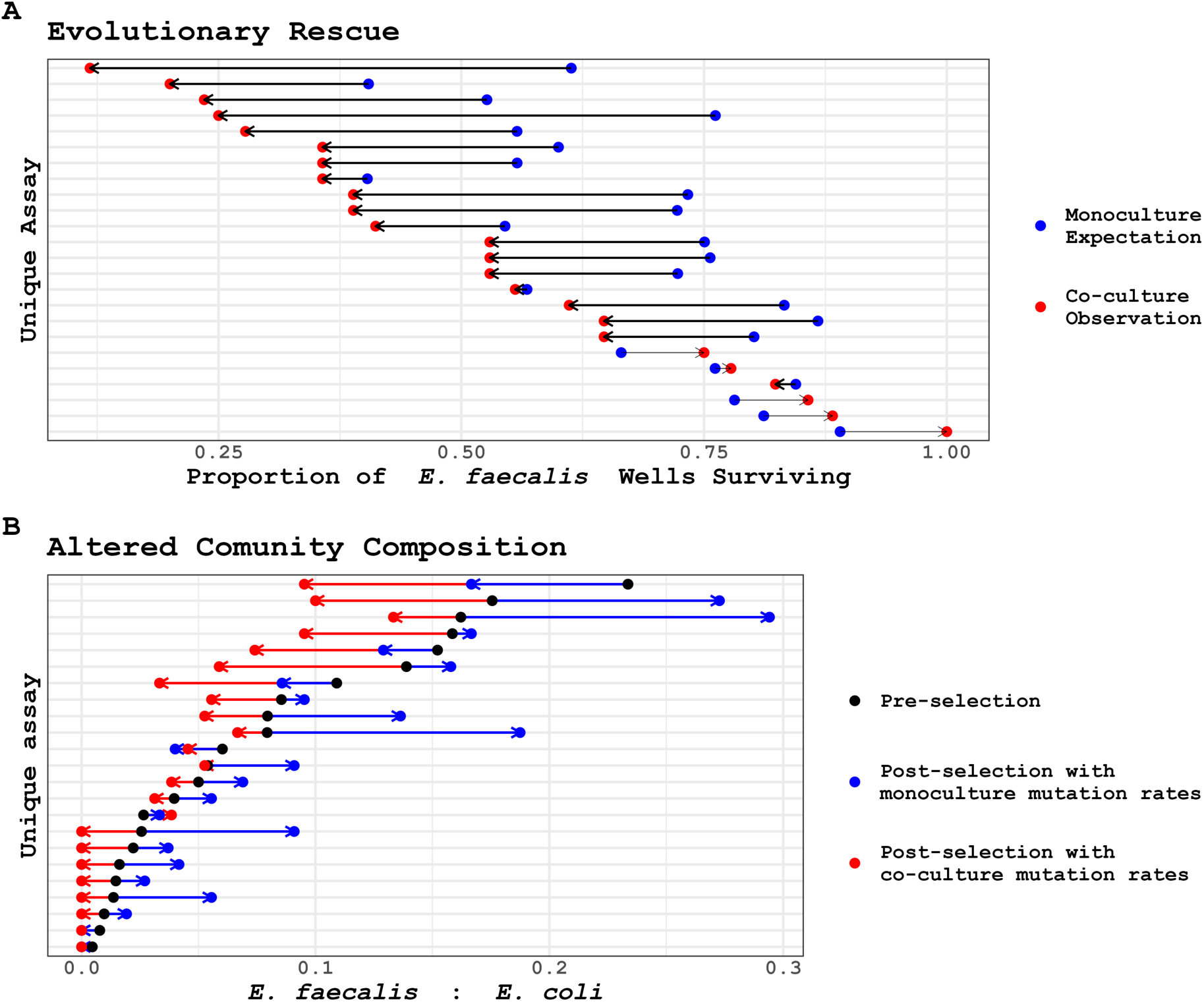
The antimutagenic effect of *E. coli* on *E. faecalis* reduces the rates of evolutionary rescue (A) in *E. faecalis* and also reduces the *E. faecalis*:*E. coli* ratio following antibiotic selection (B). A) Each pair of points shows a single co-culture fluctuation assay. Red points show observed proportion of wells with >=1 surviving *E. faecalis* colony (N_wells_ per assay = 14-18). Blue points show the expected proportion of wells with >=1 surviving *E. faecalis* colony across 10,000 simulated parallel cultures using monoculture mutation rates as fitted in Figure 3. B) Each set of 3 connected points shows a single co-culture fluctuation assay. In black the experimentally measured final ratio of *E. faecalis*:*E. coli* prior to selection is shown. Red and blue points show the expected *E. faecalis*:*E. coli* ratio across 10,000 simulations for each species using co-culture or monoculture mutation rates respectively as fitted in Figure 3. For each red or blue point 10,000 simulated parallel cultures are carried out for each species and the median values for both species used to estimate the average expected *E. faecalis*:*E. coli* ratio. Fitted monoculture and co-culture mutation counts are shown in Figure S7.

In co-cultures of *E. coli* + *E. faecalis* we find that each population density doubling of the weaker ROS degrader, here *E. faecalis*, is associated with a 1.2-fold increase in the mutation rate of the stronger ROS degrader, here *E. coli* (95% CI: 1.1 - 1.4, *t* ratio_8_ = 3.2, *P* = 0.025, Figure 4D, Statistical Model 4). However, there is no significant effect of *E. coli* density on the mutation rate of *E. faecalis* in co-culture (t ratio_8_ = 0.42, *P* = 0.69, Figure 4D, Statistical Model 4). Thus, as with *P. aeruginosa* + *E. coli*, we observe a quantitative increase in mutation rates for the stronger ROS degrader (*P. aeruginosa* > *E. coli* > *E. faecalis*) as the density of the weaker ROS degrader is increased (Figure 4), but no similar effect of the stronger degrader on the weaker, where the mutation rate is simply qualitatively decreased by co-culture (Figure 3). In both systems, the failure to find a quantitative effect of the stronger degrader on the weaker could be due to low statistical power. During *E. coli* + *E. faecalis* co-culture experiments *E. faecalis* density varies by 59-fold while *E. coli* density varies by only 2.7-fold (Figure 4C), limiting our power to detect an effect of *E. coli* density on *E. faecalis* mutation rates. Similarly, during *P. aeruginosa* + *E. coli* cocultures density-associated effects of *E. coli* are better captured than those of *P. aeruginosa* as *E. coli* densities vary by 44-fold in this dataset while *P. aeruginosa* densities vary by only 8.9-fold (Figure 4A). These results demonstrate that both the presence (Figure 3) and density (Figure 4) of a co-cultured species can alter microbial mutation rates by similar magnitudes, for example while the presence of *P. aeruginosa* decreases *E. coli* mutation rates by 1.8-fold a doubling in the population density of *E. coli* increases the mutation rate of *P. aeruginosa* by 1.4-fold and a quadrupling of population density increases mutation rates by 2-fold.

### Cross-species mutational effects complete an eco-evo feedback by altering rates of evolutionary rescue

Finally, while we have demonstrated and dissected the ecological effect of the biotic environment on an evolutionary mechanism (mutation rate), a full eco-evo feedback needs to demonstrate that this evolutionary mechanism in turn affects the ecology of the system. This is tested by the experiments already reported for estimating the mutation rate. These experiments (fluctuation assays: Figure 3, Figure 4) involve modifying the environment for each of the 4,850 experimental cultures used. Each culture was confronted with a lethal stress (the antibiotic rifampicin) and the extent of evolutionary rescue assayed. The resulting populations differ evolutionarily in their genotypes (having been selected for antibiotic resistance at the *rpoB* locus), but the resulting co-cultures also potentially differ in their ecological interactions through changes in the relative abundance of the two species. At this point we consider the potentially confounding effect of differential selection for or against rifampicin resistance (rifR) in mono vs co-culture conditions. Should coculture conditions select more strongly against rifR then both mutation rates and rates of evolutionary rescue may be underestimated. Fortunately, it is possible to coestimate both the expected number of mutational events (*m*) and the fitness effects of resistance (*ρ*^-1^) from mutant colony counts and so this can be tested for. In both systems,while costs of resistance vary among species, we find no significant or substantial effects of co-culture on the fitness effects of resistance for any species (Figure S4).

We can generate expectations for the effects of evolutionary rescue on the community composition in co-culture assays based on monoculture mutation rates (Figure 3). Figure 5A shows that the observed rates of evolutionary rescue for *E. faecalis* populations confronted with lethal rifampicin concentrations, following growth in co-culture with *E. coli*, are substantially lower than expected given monoculture mutation rates (average difference between monoculture expectations and co-culture observations =-16.6%, t_23_ =-4.7, *P* = 9.30 × 10^−5^). To further illustrate the effect of cross-species mutational effects on community composition we simulated the surviving ratio of *E. faecalis*:*E. coli* in each assay following rifampicin selection. Figure 5B shows these simulated results based on the fitted mutation rates presented in Figure 3 and Statistical Model 2. In these populations, where *E. faecalis* is initially in a minority, its higher average mutation rate than *E. coli* (Figure 3) is typically predicted to provide the genetic variation required for evolutionary rescue at a higher rate, resulting in a higher *E. faecalis*:*E. coli* ratios after antibiotic than before (i.e. most blue points in Figure 5B are higher than the black points, average difference between initial ratio and expectation under monoculture mutation rates = +0.0232, t_22_ = 2.5, *P* = 2.25 × 10^−2^). However, we can see that the strong negative effect of *E. coli* on the mutation rate (Figure 3), the only factor that differs in generating the expectations for mono-and co-cultures plotted in Figure 5B, consistently means that *E. faecalis* is instead less likely to experience evolutionary rescue (i.e. most red points in Figure 5B are lower than the black points, average difference between initial ratio and expectation under co-culture mutation rates =-0.0325, t_22_ =-4.4, *P* = 2.43 × 10^−4^). Thus, the post-selection communities have a substantially lower *E. faecalis*:*E. coli* ratio than expected, not because of selective effects but, because of the effects of co-culture on mutation rate. Knowing this effect of co-culture on mutation rate *reverses* the expected direction of community composition change following evolutionary rescue for 16 out of 23 co-culture assays (Figure 5B). We repeated this analysis for the *P. aeruginosa* + *E. coli* system finding a similar pattern in which the probability of evolutionary rescue and proportion of the post-selection population made up by *E. coli* were both reduced as a result of mutation rate suppression by *P. aeruginosa* (Figure S5, Figure S6).

## Discussion

We find community composition to shape bacterial mutation rates, which in turn, following a sudden stress and evolutionary rescue, shape community composition. This constitutes a novel form of eco-evolutionary feedback. Across both co-culture systems (*P. aeruginosa* + *E. coli* and *E. coli* + *E. faecalis*), mutation rates differed between mono-and co-culture conditions (Figure 3). In each system, mutation rates of the weaker ROS degrader, while higher to start with, were reduced in co-culture. Mutation rates were further influenced by the density of the co-resident species, with cross-species density effects observed in both systems (Figure 4). While mutation rates in monoculture decrease at higher population densities [8], this negative relationship between mutation rate and population density does not extend simply to cross-species density effects. Instead, the direction and magnitude of the mutational response to a co-resident species depends on the specific species pairing. Together, these results show that mutation rates are influenced by microbial community composition, rather than being determined solely by species identity. This community effect on mutation rate alters the availability of genetic variants in the constituent populations thereby altering community composition following environmental change, closing an eco-evolutionary feedback loop.

In addition to the mutation rate effects demonstrated here, mutation supply, the generation of genetic variation on which natural selection acts, will also be influenced by ecological context via effects on population size [71]. Changes to mutation supply may alter both evolutionary trajectories and subsequent community composition by modifying the generation of genetic variation. It is therefore possible for cross-species effects on mutation supply to alter the chances of coexistence and competitive exclusion in microbial communities.

### Mechanisms of cross-species mutational effects

This study allows us to explore mechanisms underlying cross-species mutational effects. One possible explanation is that a stronger ROS degrader removes mutagenic ROS from a shared environment, thereby reducing DNA damage and mutation rates in the co-cultured species. This hypothesis is supported by previous work showing that mutation rates in a ROS-degradation deficient strain of *E. coli* decreased as the density of co-resident wildtype *E. coli* increased [24]. However, the mutagenic effect of *E. coli* on *P. aeruginosa* (Figure 4A&B) is not explained by this mechanism as *E. coli* does not excrete any significant amount of hydrogen peroxide [72]. It is also possible that co-culturing may alter ROS concentrations indirectly by altering oxygen availability [73]. Further manipulation and direct measurement of the chemical environment during mutation rate assays, with both monoculture and co-culture systems, is required to uncover the mechanisms of within and across-species mutational effects.

### Methods for measuring cross-species mutational effects

This study also highlights important challenges to be addressed as we expand our understanding of mutation supply to complex communities and non-model microbes. An important challenge in estimating mutation rates arises in organisms with variable effective ploidy, such as *E. faecalis*, which forms diplococci and short chains (Figure 2). As a result, colonies may originate from more than one founder cell [74–76], introducing uncertainty into both mutant counts and population-size estimates derived from colony forming unit (CFU) counts. Highly variable effective ploidy (genomes per observational unit) is common among bacterial species that form filaments or multicellular aggregates, including *E. faecalis* [77] and is also observed in other biological systems including polynucleate fungal hyphae [78], cancers with frequent chromosome gain and loss [79] and plants with tissue-specific ploidy variation [80]. In such systems, ploidy is more accurately described as a distribution rather than a single value. Unfortunately, despite the known importance of ploidy in mutation rate estimation [81–83], approaches that explicitly account for ploidy variation within a population are not currently available. Developing approaches that account for variation in chain length would allow fluctuation assay analyses to be extended to aggregate-forming bacteria, such as *Staphylococcus aureus* and *Acinetobacter baumannii*, and would enable re-analysis of datasets where aggregation may have biased mutation-rate estimates (e.g. [21]).

Here we estimated mutation rates using the fluctuation assay [69, 70] however, given the challenges in meeting the assumptions of this assay, alternative approaches should be considered. The popular mutation accumulation assay [69] has similar drawbacks to the fluctuation assay, as this approach relies on true single-cell bottlenecks and is therefore not well suited to organisms that grow as multicellular aggregates. Deep population sequencing, when selection and genetic drift are minimised, may provide the most promising approach to estimating mutation rates in non-model species and complex communities [84–89]. By detecting mutations directly from population DNA rather than inferring them from colony counts, this method avoids biases introduced by aggregation however it will remain important to account for mortality.

Bacteria frequently harm or kill competitors through diverse mechanisms of interference competition, including chemical, mechanical and biological weapons [90]. Mortality is therefore an important consideration when estimating mutation rates, as fluctuation assays typically assume minimal death during population growth. This problem has been highlighted in studies of antibiotic stress, where cell death during treatment led to inflated mutation-rate estimates when mortality was not accounted for [91]. We therefore hypothesised that *E. coli* mortality may increase when grown in co-culture with *P. aeruginosa* due to *P. aeruginosa*’s extensive bacterial warfare mechanisms [90]. However, we observed no significant increase in cell death in co-culture compared with monoculture conditions (Figure S8, Figure S9). Nonetheless, propidium iodide staining only detects intact cells with compromised membranes, meaning that cells which fully lyse during the fluctuation assay would not be detected. This is particularly relevant if *P. aeruginosa* employs lytic defence mechanisms, such as those delivered by the Type VI secretion system (T6SS), which could result in undetected *E. coli* death [92]. Mortality therefore remains an important consideration as studies of mutational processes expand into complex community contexts.

## Summary

In summary, our findings show that mutation rates are influenced by community composition and vice versa. Ecological context is typically considered to shape evolution primarily through changes in selective pressures, yet our results show that interactions between species can also alter the supply of genetic variation on which selection acts. Mutational spectra are also known to vary with environmental conditions and ecological niche [95], so interspecies interactions may influence not only the rate at which mutations arise but also the types of mutations that are available to selection. Those mutational spectra, as with the rate effects explored here, may similarly feed back into the community’s ecological interactions and evolutionary trajectory over longer time-periods. This emphasises the possibility that the mutation rates and mutational spectra of polymicrobial infections may differ from those measured in monoculture, with the direction of the effect depending on the particular species involved, making eco-evolutionary prediction very complex. Nonetheless, should ROS phenotypes prove generally predictive of these effects, in the way that they are for the species pairs studied here, mutation rates in a community context may remain predictable despite the complex and dynamic nature of microbial mutagenesis.

## Supporting information

Graphical Abstract

## Acknowledgements

The authors would like to María Rebolleda Gómez and Laurinne J Balstad for insightful discussion. We also thank Johanna M. Schwingel for *P*. *aeruginosa* PAO1, Mato Lagator for *E. faecalis* ATCC19433 and Mike Bottery & Mike Brockhurst for *E. coli* MV1463. Thanks finally to MicrobesNG (http://www.microbesng.com) for their genome sequencing services.

## Data Availability

All code and data are available on the OSF: https://osf.io/c8mda

Raw data files are as follows:

DataGuide.xlsx describes all variables in the data tables listed here

Data1_FA_Efaecalis_Ecoli_raw_data – Raw fluctuation-assay data for *E. coli* and *E. faecalis*, including monoculture and coculture conditions.

Data2_All_PA01_fluctuation_assays-Raw fluctuation-assay data for *E. coli* and *P. aeruginosa* mutation rates across all experiments, including monoculture and coculture.

Data3_All_Ef_Ec_FA_processed-Processed fluctuation-assay data for *E. coli* and *E. faecalis*, including monoculture and coculture conditions, mutation rate estimates and fitness estimates.

Data4_All_Pa_Ec_FA_processed-Processed fluctuation-assay data for *E. coli* and *P. aeruginosa* in monoculture and coculture conditions, mutation rate estimates and fitness estimates.

Data5_Chain_lengths_2026-*E. faecalis* chain-length measurements from monoculture and *E. coli* coculture conditions, manually counted from microscopy images (annotated images can be found in the OSF repository).

Data6_RawData_FlowCytometry-Raw flow-cytometry data used to assess cell viability in *E. coli* and *P. aeruginosa* during fluctuation assays in monoculture and coculture conditions.

Data7_EFrescue.csv - Observed and monoculture model-predicted survival probabilities for *E. faecalis* cocultures faced with lethal rifampicin selection used to generate Figure 5.

Data8_EF_EC_rescue.csv - Data used to estimate changes in *E. coli*-*E. faecalis* community composition following antibiotic selection using monoculture and coculture mutation rates as shown in Figure 5B

Data9_Figure_PArescue-Observed and expected *E. coli* survival data used to assess *E. coli* evolutionary rescue in *E. coli*-*P. aeruginosa* cocultures, as shown in Figure S5A.

Data10_Figure_PArescue-Data used to estimate changes in *E. coli*-*P. aeruginosa* community composition following antibiotic selection using monoculture and coculture mutation rates as shown in Figure S5B.

Data11_WGS - Whole-genome sequencing of *E. coli* and *E. faecalis* wildtype stocks to identify differences from type strain sequence.

Data12_Media_Table - Ingredients for all media used.

