## Supplement for "Cohabiting microbes can impact one another’s mutation rates creating a mutational eco-evo feedback loop"

### Supplementary Figures, Tables & Text

#### Supplementary text 1: Fluctuation assays for cocultures and aggregate forming bacterial species

##### Multicellular aggregates in monoculture and co-culture conditions

Many bacterial species form multicellular aggregates either by the aggregation of independent cells or through the continued attachment of mother and daughter cells, violating the assumption of a fluctuation assay that plated colonies are formed by single cells. *P. aeruginosa* has been observed to form multicellular aggregates through cell collision with extracellular polymeric substrates [96, 97] while *E. faecalis* is known to form diplococci and short chains through the continued association of mother and daughter cells following division [98].

To assess any multicellular aggregates in our co-culture systems we imaged monoculture and co-culture populations from both co-culture systems using phase contrast and fluorescence microscopy. All images were taken in stationary phase cultures, reflecting the timepoint of plating during our mutation rate assays. In *P. aeruginosa* + *E. coli* co-cultures we did not observe any multicellular aggregates of either species under any condition (Figure 2). Although aggregates of *P. aeruginosa* are frequently observed in liquid culture these disperse during nutrient-limitation explaining the absence of aggregates in our stationary phase cultures [97].

In the *E. coli* + *E. faecalis* system we observed multicellular chains of varying length for *E. faecalis* when grown both in monoculture and co-culture (Figure 2). The growth of colonies on agar from a chain, as opposed to a single cell, is likely to lead to an overestimate of the per genome mutation rate. This is due to the multiple genomes in a chain, any of which, if spontaneously resistant, would form a resistant colony, despite a colony forming unit being counted only as a single opportunity for mutation. This is analogous to a dominant trait under increased ploidy [83]. Our hypothesis that *E. faecalis* mutation rates will be decreased during co-culture with the stronger ROS degrading species *E. coli* may be confounded if chains are substantially longer in monoculture than co-culture conditions. We do not find evidence for this confounding effect as *E. faecalis* chains are marginally shorter in monoculture than co-culture conditions (mean length in monoculture = 3.9, 95% CI: 3.3 - 4.5, mean length in co-culture = 4.9, 95% CI: 4.4 - 5.5, $z = -2.5$, $P = 0.012$, Statistical Model 5, Figure S10 and see supplementary microscopy files). We therefore conclude that differences in chain length are unlikely to explain any observation of *E. faecalis* mutation rates being decreased during co-culture in this system. However, this study does highlight the pressing need for methods allowing the estimation of microbial mutation rates in species for which a distribution of aggregate sizes exists and may vary systematically between treatments.

##### Mortality in monoculture and co-culture conditions

The assumption of minimal mortality during fluctuation assays may also be violated in complex environments such as co-culture. Mortality leads to an underestimation in the number of divisions in which mutations may have occurred, and any mortality will therefore lead to the overestimation of mutation rates [91]. We hypothesised that *E. coli* cell death may increase in co-culture with *P. aeruginosa*, due to *P. aeruginosa*'s extensive bacterial warfare mechanisms including Type VI secretion system and S-type pyocin toxins [90]. To assess mortality in *P. aeruginosa* + *E. coli* co-cultures we used flow cytometry with viability staining using propidium iodide (PI). We find minimal mortality (3-5%) in both species at varying population densities in both monoculture and co-culture conditions (Figure S8), validating our PI stain by inducing mortality with methanol exposure or heat treatment (Figure S9). These results suggest that the assumption of minimal mortality in co-cultures of *P. aeruginosa* + *E. coli* is met, however, PI staining will only reveal intact dead cells and so it remains possible that mortality leading to rapid disintegration of the cell may take place. The possibility of increased mortality rates is an important consideration for future studies on mutation rates in mixed species communities and has frequently confounded studies on the mutational effects of antibiotic exposure [91].

#### Supplementary figures

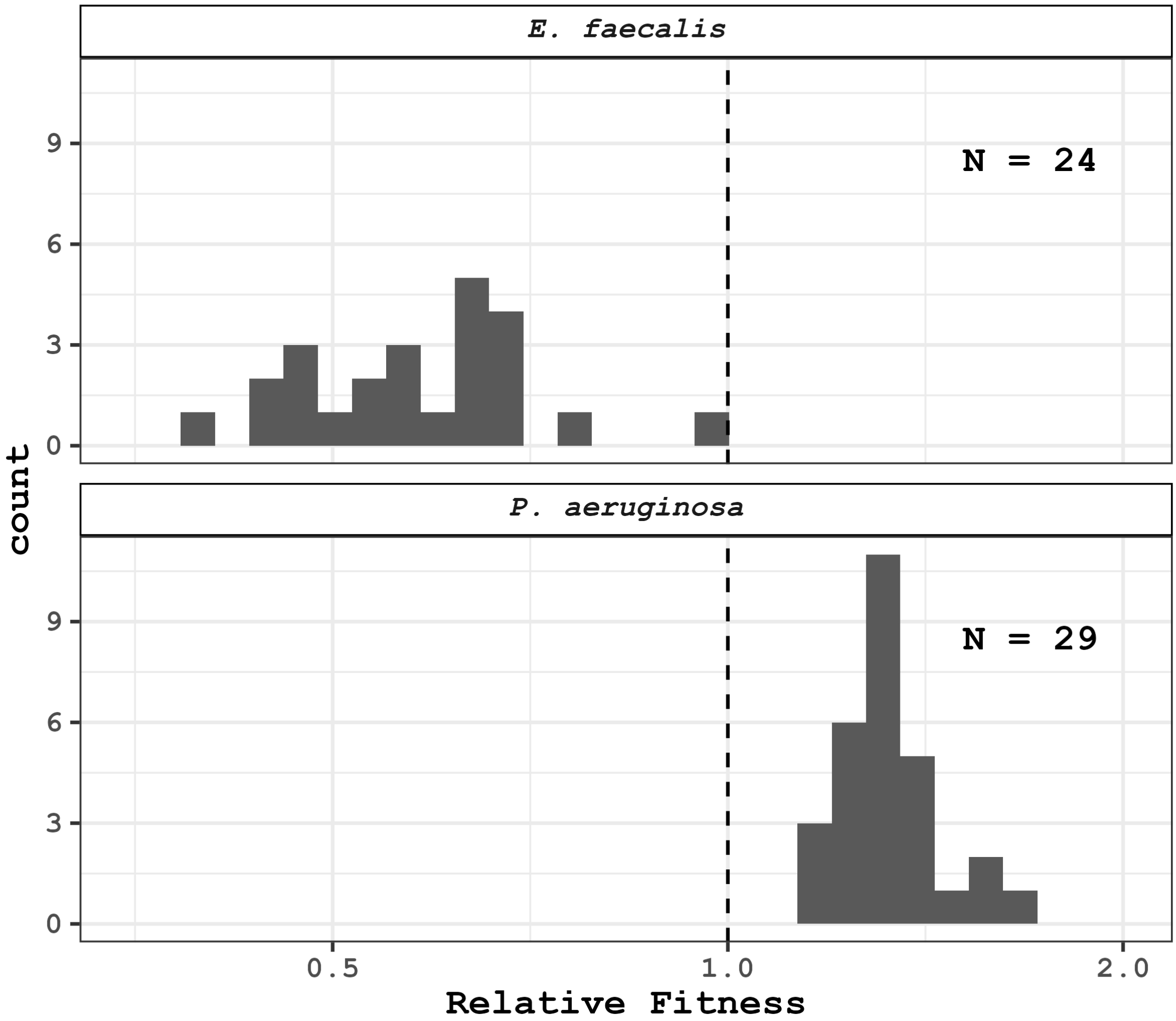


Figure S1: **Relative fitness of *P. aeruginosa* (top) and *E. faecalis* (bottom) in reference to *E. coli*.** Relative fitness is an estimate of the number of generations completed by the given species relative to each generation completed by *E. coli* during growth in co-culture. Note log scale x-axis as relative fitness is a ratio.

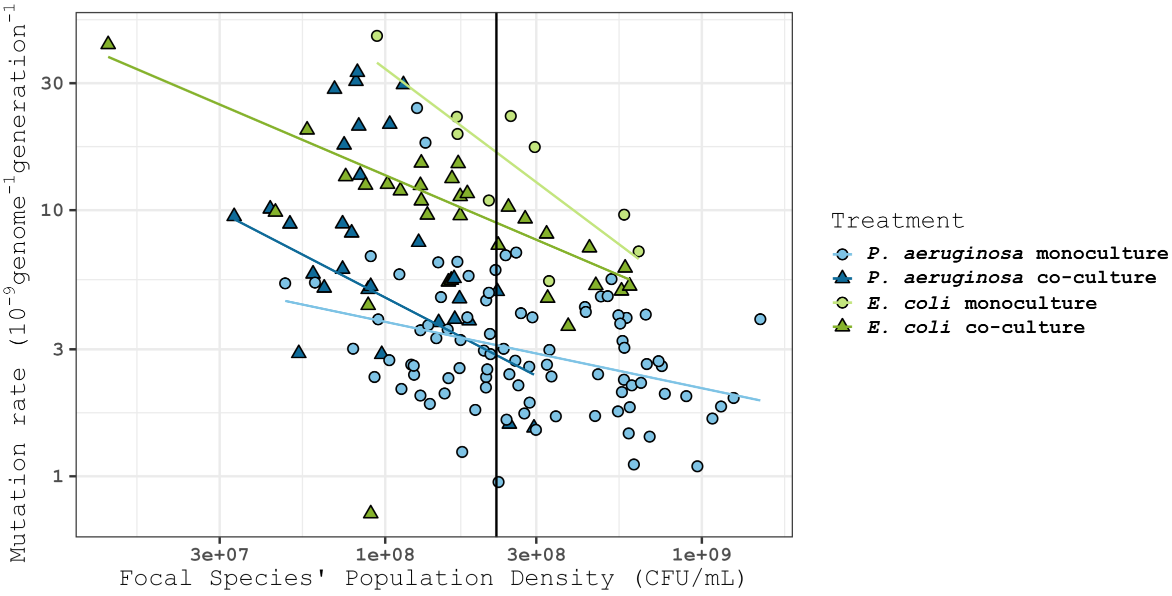


Figure S2: **Mutation rate estimates for *P. aeruginosa* & *E. coli* in monoculture and co-culture conditions.** These raw data underlie the estimates shown in Figure 3 . Blue shapes show *P. aeruginosa* and green shapes show *E. coli*, pale circles show monocultures and dark triangles show co-cultures. Sample sizes can be found in the caption of Figure 3 . Lines are fitted by statistical model 1.

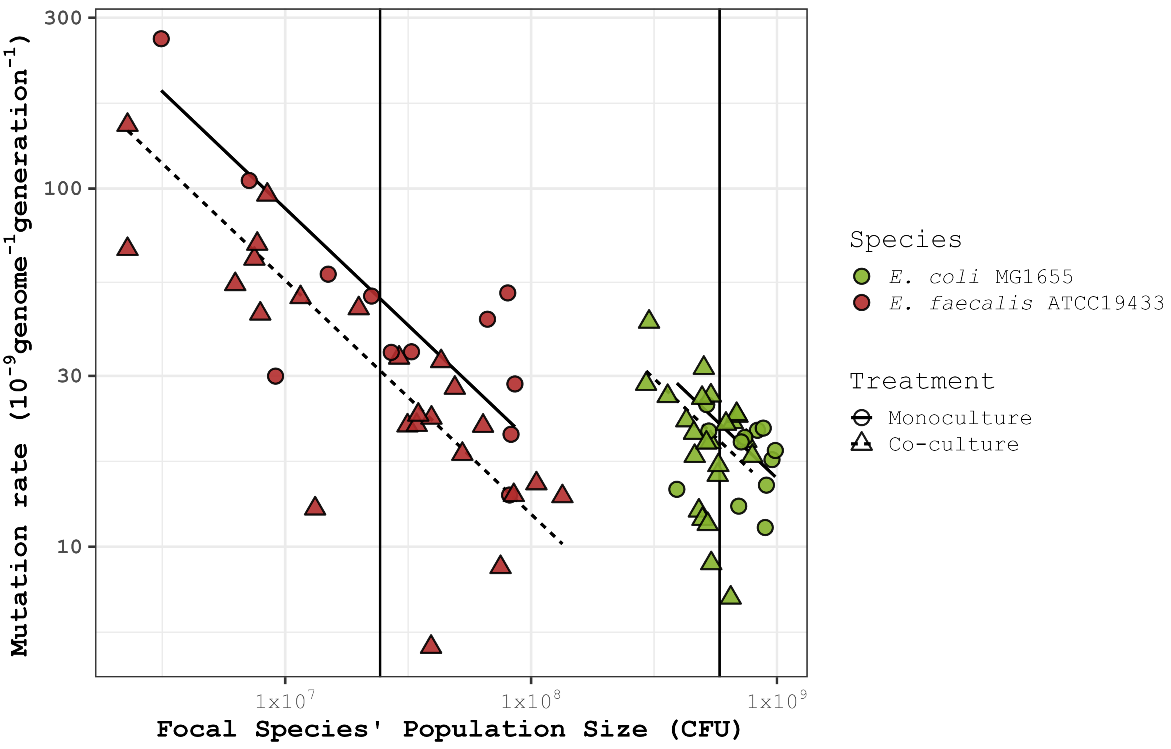


Figure S3: **Mutation rate estimates of *E. coli* & E. faecalis in monoculture and co-culture conditions.** These raw data underlie the estimates shown in Figure 3 . Red shapes show *E. faecalis* and Green shapes show *E. coli*, circles and solid lines show monocultures and triangles and dashed lines show co-cultures. Sample sizes can be found in the caption of Figure 3 . Lines are fitted by statistical model 2. We establish here that *E. faecalis* shows elevated mutation rates at low population densities i.e density-associated mutation rate plasticity (DAMP). This result is unlikely to be an artifact of cellular chain formation for the following reasons. In *E. faecalis*, colony growth on rifampicin selective media is likely to be permitted even when only a subset of a chain of cells carry a rifR mutation, effectively generating a dominant phenotype across the multiple genomes of the chain (dominance effects are discussed in [83]). Simulations assuming fixed ploidy show that when the selected trait is dominant and ploidy is >1, mutation rates are overestimated and the degree of overestimation is greatest when mutation rates are low [83]. Applying this expectation to our data suggests that multicellular chaining would lead to an underestimation, rather than inflation, of the DAMP slope in *E. faecalis* (Figure S3). We therefore conclude that *E. faecalis* exhibits DAMP and that this result is not confounded by chain formation. However, these simulations remain somewhat simplistic as they assume constant ploidy and population size [83], whereas *E. faecalis* displays variable chain lengths under our experimental conditions (Figure 2, Figure S10).

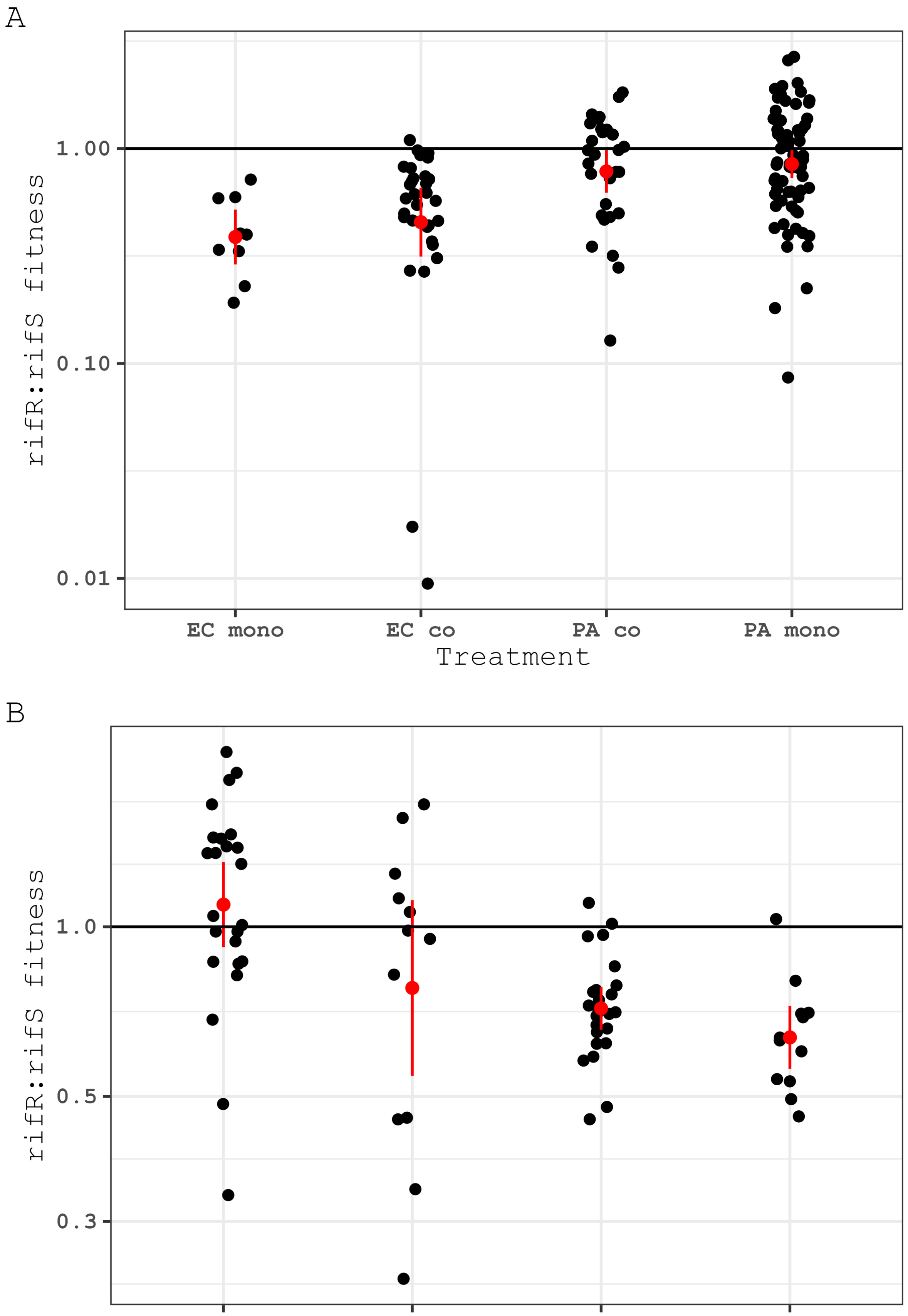

A
1.00
0.10
0.01
rifR:rifS fitness
EC mono
EC co
PA co
PA mono
Treatment
B
1.0
0.5
0.3
rifR:rifS fitness
EF co
EF mono
EC co
EC mono
Treatment

Figure S4: Fitness effects of rifampicin resistance (rifR) relative to the susceptible population (rifS) as co-estimated with mutant counts using the R package flan is shown on the y-axis. Points above the horizontal black line indicate a fitness benefit of resistance during growth in minimal media while points below the line indicate a cost of resistance. Raw data is shown in black with the mean ± 2 standard errors for each group shown in red. In panel A data is shown for the *E. coli* (EC) + *E. faecalis* (EF) system and in panel B data is shown for the *P. aeruginosa* (PA) + E. coli* (EC) system. Mono = monoculture, co = co-culture. Note log scale y-axes. For each pair a Welch Two Sample t-test was carried out with no significant difference found in any case: Ec mono vs Ec co (+ Pa) t=0.675 , df=32 , *P*=0.505 ; Pa mono vs Pa co (+ Ec) t=0.581 , df=53.4 , *P*=0.564 ; Ef mono vs Ef co (+ Ec) t=-1.71 , df=16.3 , *P*=0.106 ; Ec mono vs Ec co (+ Ef) t=1.52 , df=21.3 , *P*=0.142.

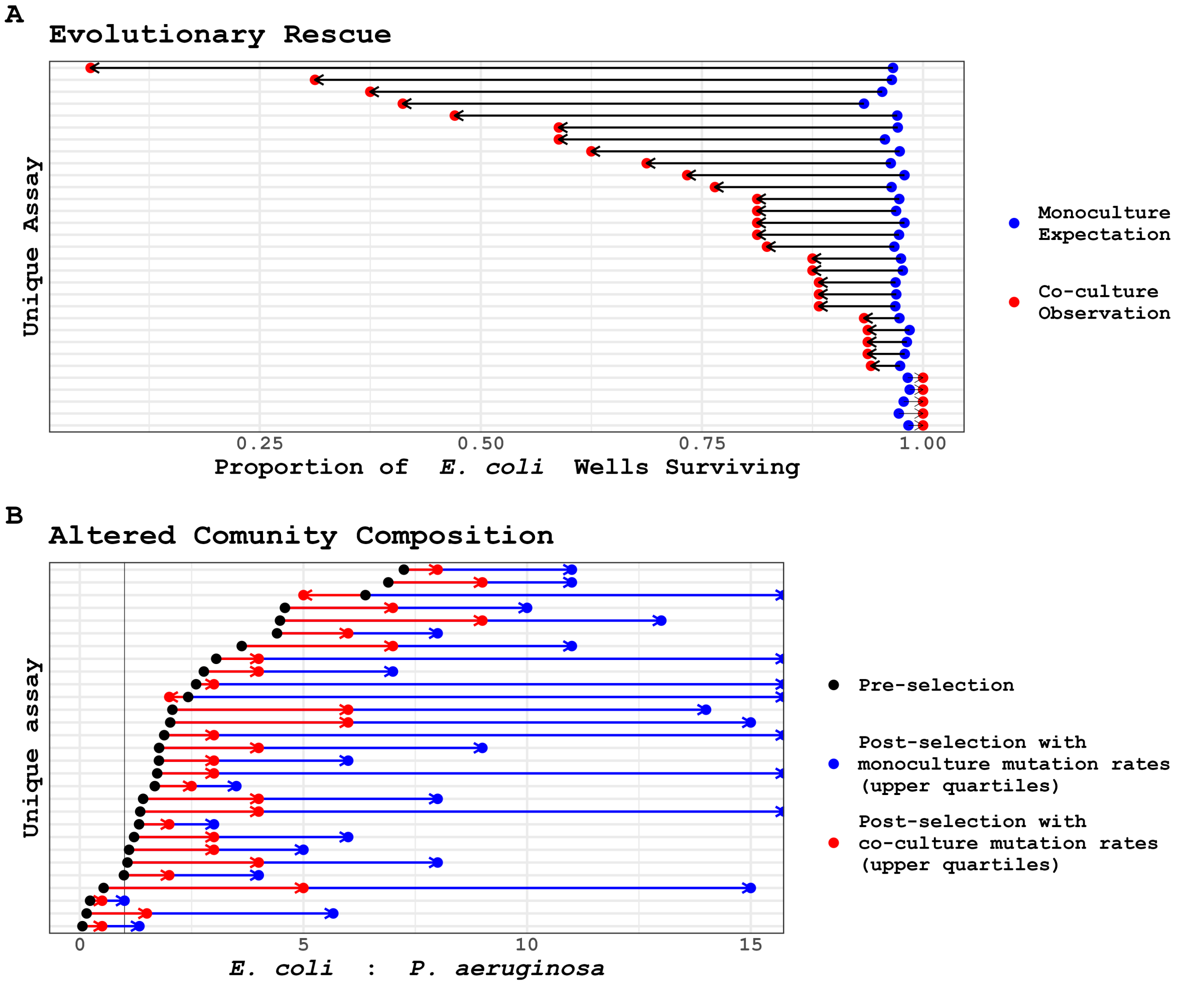


Figure S5: **The antimutagenic effect of *P. aeruginosa* on *E. coli* reduces the rates of evolutionary rescue in *E. coli* (A) and reduces the *E. coli*:*P. aeruginosa* ratio following antibiotic selection (B).** A) Each pair of points shows a single co-culture fluctuation assay. Red points show observed proportion of wells with >=1 surviving *E. coli* colony (N wells = 14-17). Blue points show the expected proportion of wells with >=1 surviving *E. coli* colony across 10,000 simulations using monoculture mutation rates as fitted in Figure 3. B) Each set of 3 connected points shows a single co-culture fluctuation assay. In black the final ratio of *E. coli*:*P. aeruginosa* prior to selection is shown. Red and blue points show the expected *E. coli*:*P. aeruginosa* ratio across 10,000 simulations for each species using co-culture or monoculture mutation rates respectively as fitted in Figure 3. For each red or blue point 10,000 simulations are carried out for each species and the upper quartile values for both species used to estimate the average *E. coli*:*P. aeruginosa* ratio. Upper quartiles are used as the median number of surviving *P. aeruginosa* colonies is frequently 0 leading to many infinite estimates. Fitted monoculture and co-culture mutation counts are shown in Figure S6.

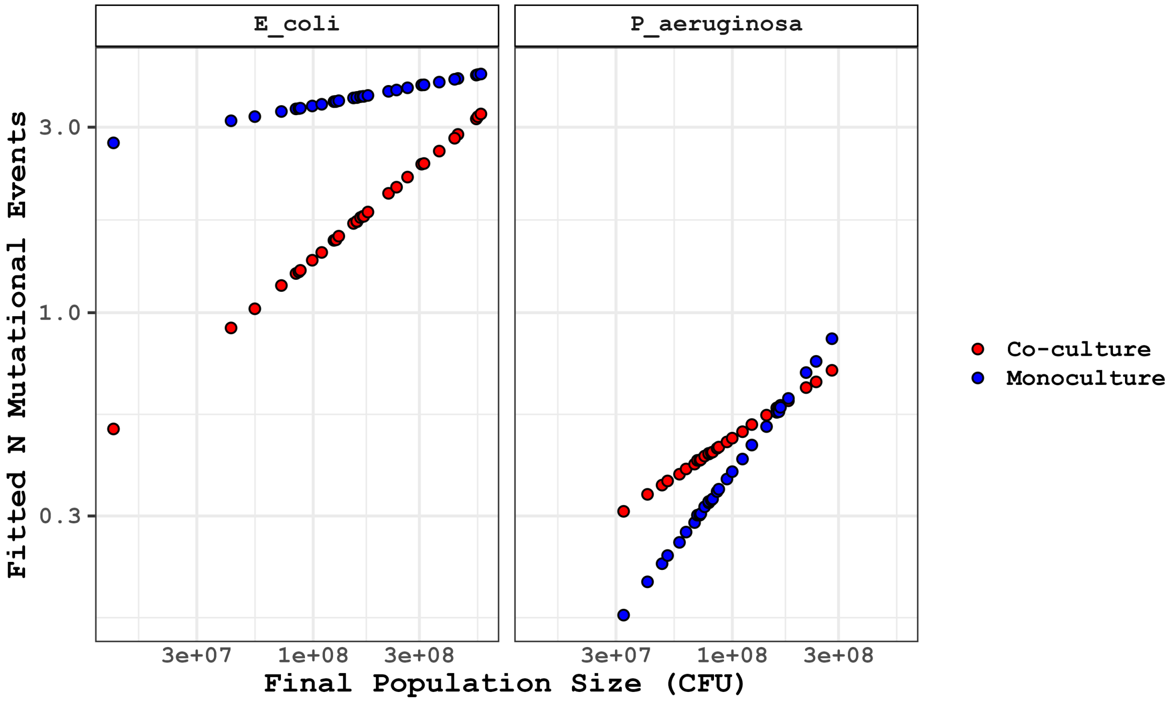


Figure S6: **Fitted mutation event (m) estimates for *E. coli* (left) and *P. aeruginosa* (right)** using statistical model 1 as shown in Figure 3. Final population sizes are taken from experimental data relating to each co-culture fluctuation assay, mutation event counts are estimated assuming either co-culture (red) or monoculture (blue) conditions. Note that mutation events (m) are not the same as mutant counts (mc), m =< mc as a single mutational event can produce >1 mutant colony if the event is early enough for subsequent reproduction prior to selective plating. x- and y-axes are plotted on a log scale.

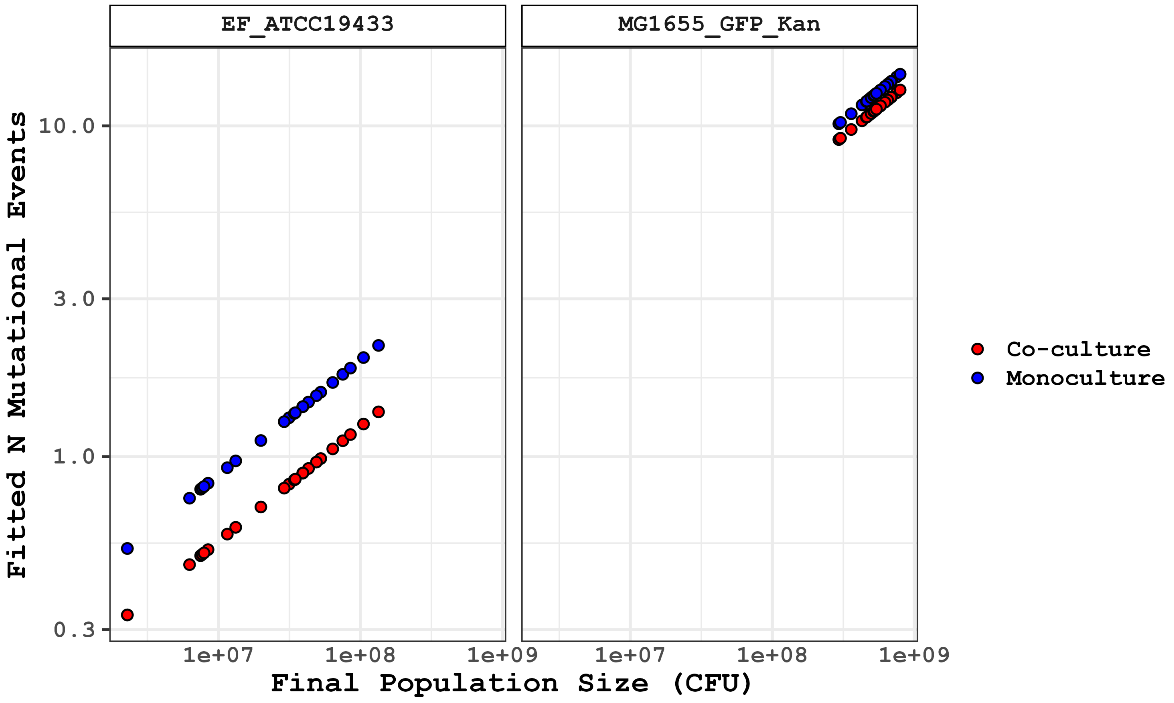


Figure S7: **Fitted mutation event (m) estimates for *E. faecalis* (left) and *E. coli* (right)** using statistical model 2 as shown in Figure 3 . Final population sizes are taken from experimental data relating to each co-culture fluctuation assay, mutation event counts are estimated assuming either co-culture (red) or monculture (blue) conditions. Note that mutation events (m) are not the same as mutant counts (mc), m =< mc as a single mutational event can produce >1 mutant colony if the event is early enough for subsequent reproduction prior to selective plating. x- and y-axes are plotted on a log scale.

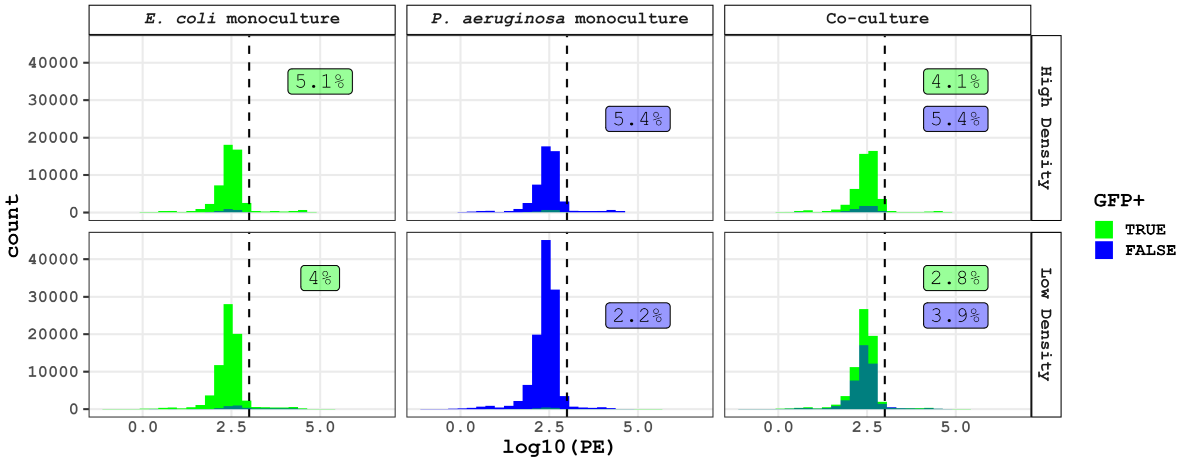


Figure S8: **Minimal mortality observed in monocultures and co-cultures of *P. aeruginosa* + *E. coli*.** Histogram of propidium iodide stain intensity with a value of log10(PE)>3 classified as dead based on positive controls for death (see Figure S9 for all raw data). In each panel the relevant mortality rates are shown for GFP+ events (green background, y = 35,000) and GFP- events (blue background, y=25,000).

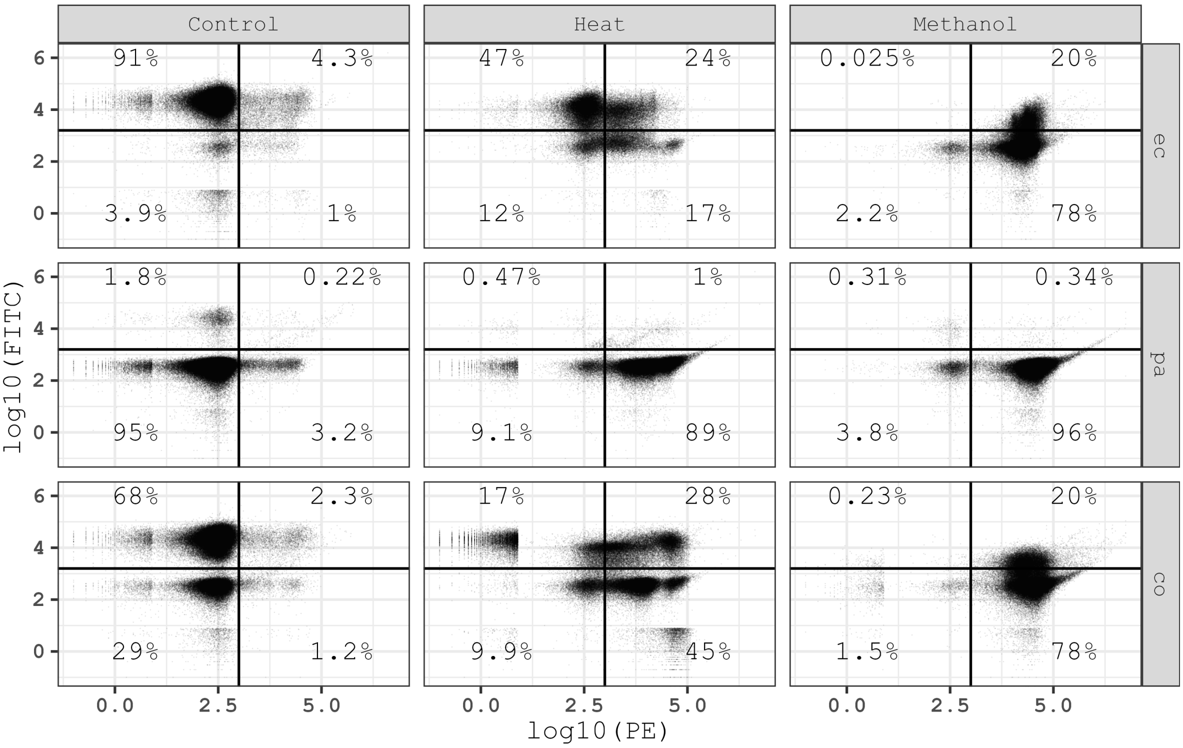


Figure S9: Individual events captured during flow cytometry of *E. coli* and *P. aeruginosa* monocultures and co-cultures. Heat and methanol treatments are used as positive controls for PI staining of dead cells. From top left clockwise quadrants can be interpreted as GFP+PI-, GFP+PI+, GFP-,PI+,GFP-PI- with PI+ being dead cells and PI- being live cells. The *E. coli* strain used here constitutively expresses GFP however this is seen to differing extents during heat and methanol treatment (see Table 1). Percentages show the proportion of events in the given condition with the quadrant phenotype.

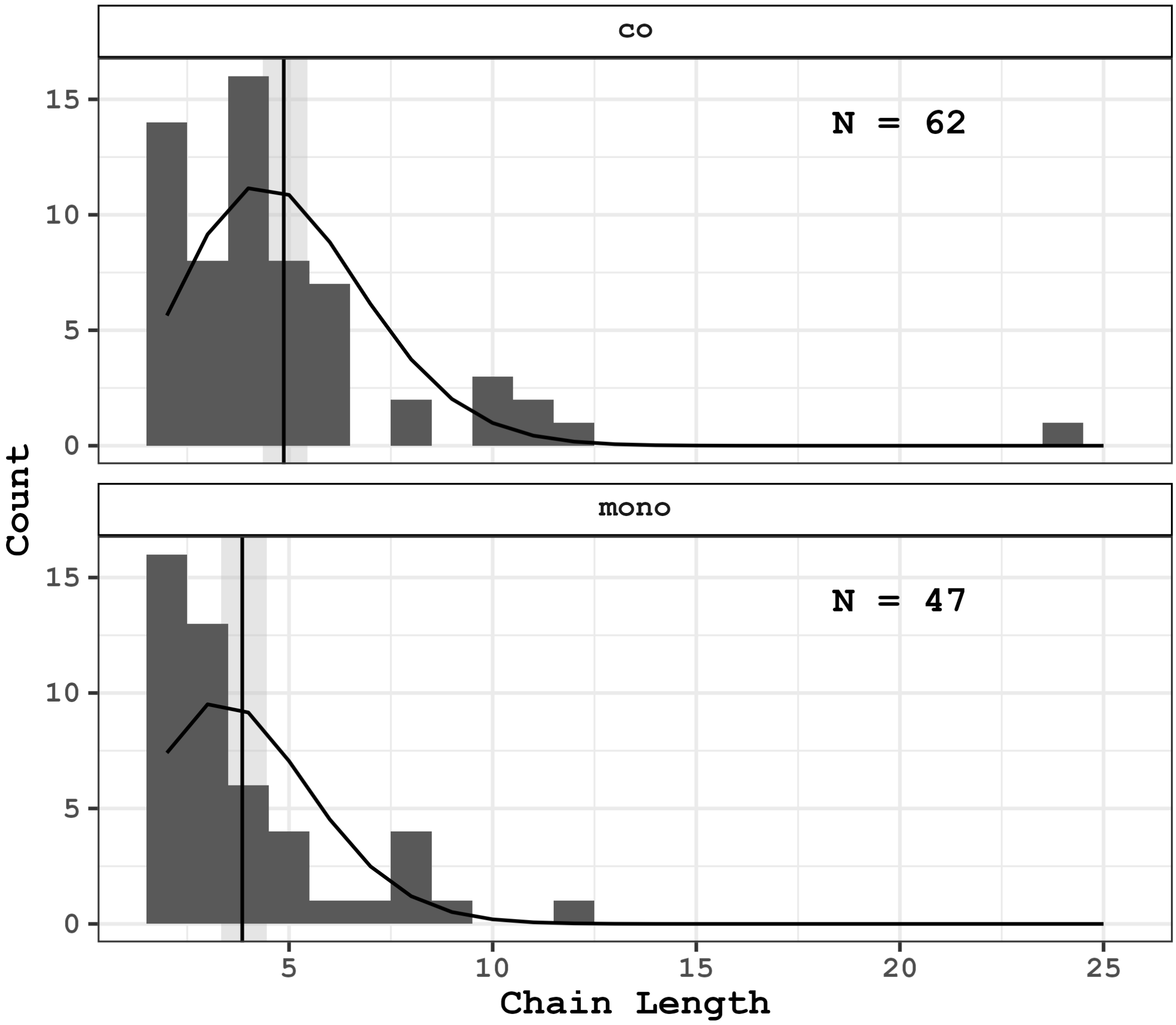


Figure S10: **Distribution of *E. faecalis* chain lengths when grown in co-culture (top) and monoculture (bottom).** A Poisson fit to this data using Statistical Model 5 is shown as a black curve. The mean chain length and 95% confidence intervals are shown by a vertical black line and shaded grey region. For both co- and monoculture all chains of length ≥2 in 3 separate microscopy images were counted, the total number of chains counted for each condition is shown in the top righthand of the plot (see Methods for details). Chains of length 1 are not included as *E. faecalis* cannot be distinguished from *E. coli* cells not expressing GFP (a significant minority of *E. coli* cells).

#### Supplementary table

Table 1: **Table S1: Summary table of all events observed by flow cytometry split by GFP+/- status.** For each treatment/glucose combination the number of biological replicates, total events, GFP- events, GFP+ events and proportion of events GFP+ are given.

| Sample | Treatment | Glucose | Replicates | Total Events | GFP- | GFP+ | ProportionGFP+ |
|---|---|---|---|---|---|---|---|
| co | Control | 1000 | 3 | 54489 | 5843 | 48646 | 89.3% |

| | | | | | | | |
|---|---|---|---|---|---|---|---|
| co | Control | 250 | 6 | 112068 | 44201 | 67867 | 60.6% |
| ec | Control | 1000 | 3 | 54801 | 2719 | 52082 | 95% |
| ec | Control | 250 | 4 | 75197 | 3654 | 71543 | 95.1% |
| pa | Control | 1000 | 3 | 54763 | 52599 | 2164 | 3.95% |
| pa | Control | 250 | 6 | 116416 | 115085 | 1331 | 1.14% |
| co | Heat | 1000 | 1 | 18162 | 3573 | 14589 | 80.3% |
| co | Heat | 250 | 4 | 89212 | 55860 | 33352 | 37.4% |
| ec | Heat | 1000 | 1 | 18312 | 4289 | 14023 | 76.6% |
| ec | Heat | 250 | 4 | 56175 | 17539 | 38636 | 68.8% |
| pa | Heat | 1000 | 1 | 18369 | 18022 | 347 | 1.89% |
| pa | Heat | 250 | 4 | 66957 | 66056 | 901 | 1.35% |
| co | Methanol | 1000 | 3 | 55486 | 40951 | 14535 | 26.2% |
| co | Methanol | 250 | 3 | 56171 | 47984 | 8187 | 14.6% |
| ec | Methanol | 1000 | 3 | 55830 | 41387 | 14443 | 25.9% |
| ec | Methanol | 250 | 3 | 58215 | 50253 | 7962 | 13.7% |
| pa | Methanol | 1000 | 3 | 55804 | 55579 | 225 | 0.403% |
| pa | Methanol | 250 | 3 | 57467 | 56958 | 509 | 0.886% |

### Statistical Appendix to Green & Wright et al

Standard diagnostic plots are included for all linear models. All models were executed in R. Although the document can be read with no reference to the code we have chosen to also include code elements for those it might benefit. Full code as a quarto document associated with this paper is available in the supplement.

#### Statistical Model 1

The model shown in figure 1 (lefthand side) and Figure S2 fits $\log_2$ mutational events ($m$) per mL against the mean-centred $\log_2$ density (Dc), allowing for differences in intercept and slope among all 4 treatments (*P. aeruginosa* in monoculture, *P. aeruginosa* in co-culture, *E. coli* in monoculture and *E. coli* in co-culture). Random effects of experimental plate (47 levels) nested within experimental block (23 levels) nested within experimenter (4 levels), each affecting the intercept, are also fitted. Differences in variance (i.e. heteroscedasticity) associated with the upper bound of estimated mutational events per mL were also included varying by identity of the co-cultured species.

Predicted values from this model shown as lines of best fit in Figure S2 are calculated by dividing predicted mutational events per mL from this model by the measured value of CFU per mL (population density); both the ‘per mL’ terms cancel and the result is in the units of mutational events per CFU, therefore giving a mutation rate.

#### Model call

```
model <- lme(log2(m_mL)~Dc*treatment, data=d2,
    random=~1|Experimentor/block/plate_ID,
    weights = varPower(form = ~m_upper_genotype | coculture_strain))
```

#### Coefficient table for model 1

| | Value | Std.Error | DF | t-value | p-value |
|---|---|---|---|---|---|
| (Intercept) | 1.890 | 0.367 | 124 | 5.150 | 9.85e-07 |
| Dc | 0.108 | 0.322 | 124 | 0.334 | 0.739 |
| treatmentMG1655_GFP_Kan_PA01 | -0.879 | 0.278 | 124 | -3.160 | 0.00196 |
| treatmentPA01_MG1655_GFP_Kan | -2.530 | 0.393 | 124 | -6.450 | 2.32e-09 |
| treatmentPA01_none | -2.410 | 0.326 | 124 | -7.380 | 2.03e-11 |
| treatmentPA01_none_LB | -0.851 | 0.548 | 23 | -1.550 | 0.134 |
| Dc:treatmentMG1655_GFP_Kan_PA01 | 0.384 | 0.329 | 124 | 1.170 | 0.245 |
| Dc:treatmentPA01_MG1655_GFP_Kan | 0.275 | 0.373 | 124 | 0.739 | 0.461 |
| Dc:treatmentPA01_none | 0.643 | 0.334 | 124 | 1.930 | 0.0562 |
| Dc:treatmentPA01_none_LB | 0.966 | 0.487 | 124 | 1.980 | 0.0497 |

#### ANOVA table for model 1

| | numDF | denDF | F-value | p-value |
|---|---|---|---|---|
| (Intercept) | 1 | 123 | 2.54 | 0.113 |
| Dc | 1 | 123 | 232.00 | <2.2e-16 |
| treatment | 4 | 123 | 37.10 | <2.2e-16 |
| Dc:treatment | 4 | 123 | 2.62 | 0.0384 |

#### Variance and Standard Deviations of model 1

| | Variance | StdDev |
|---|---|---|

| | | |
|---|---|---|
| Experimentor = | 0.1350 | 3.67e-01 |
| block = | 0.0878 | 2.96e-01 |
| plate_ID = | 0.0000 | 6.69e-05 |
| Residual | 0.6080 | 7.80e-01 |

##### Weights vary with the upper bound of *m* to the following powers

| **coculture_strain** | **varPower** |
|---|---|
| none | 0.002536917 |
| MG1655_GFP_Kan | 0.604344577 |
| PA01 | -0.690629698 |

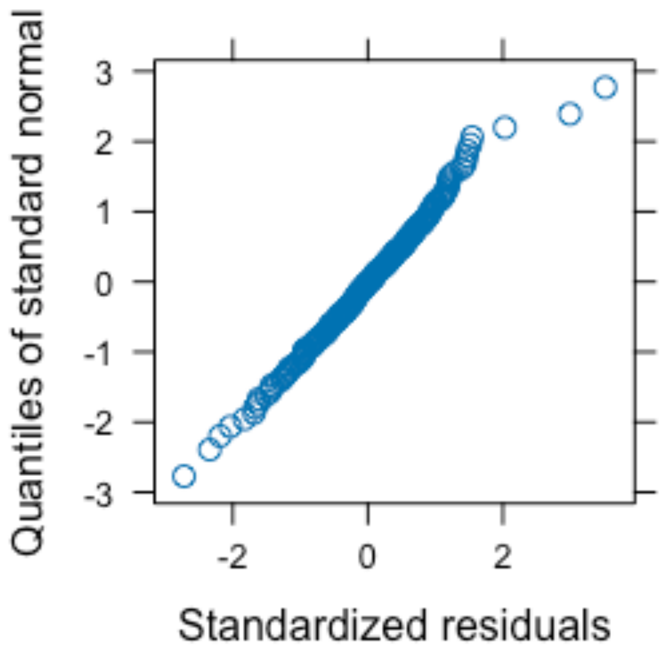


QQ Plot of Deviance Residuals for Model 1

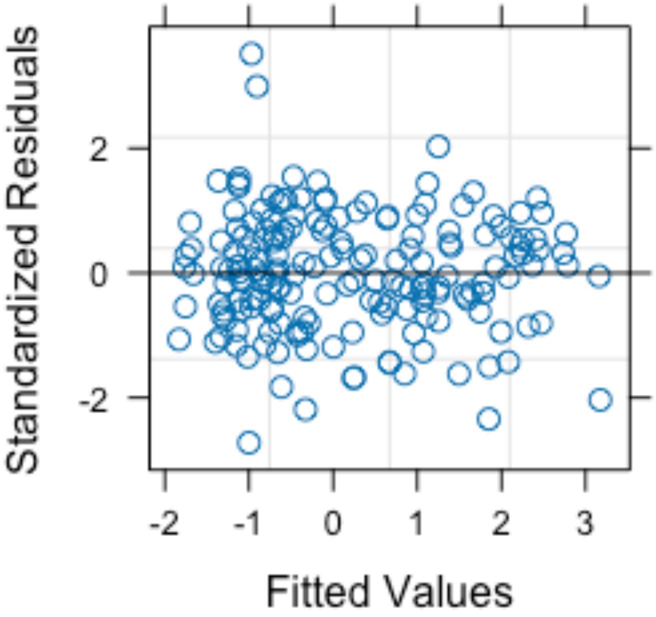


Standardized Residuals vs Fitted Values for Model 1

#### Statistical Model 2

The model shown in figure 1 (righthand side) and Figure S3 fits $\log_2$ mutational events (*m*) per culture against the mean-centred $\log_2$ final population size (Nc), allowing for differences in intercept among all 4 treatments (*E. faecalis* in monoculture, *E. faecalis* in co-culture, *E. coli* in monoculture and *E. coli* in co-culture). Random effects of experimental plate (15 levels) nested within experimental block (5 levels), each affecting the intercept, are also fitted. Differences in variance (i.e. heteroscedasticity) associated with the product of the relative fitness of the wildtype to the rifR mutants and the percentage of casamino acid supplementation were also included.

Predicted values from this model shown as lines of best fit in Figure S3 are calculated by dividing predicted mutational events per culture from this model by the measured value of CFU, therefore giving a mutation rate.

Model 2 was originally fitted allowing differences in slope among treatments however this interaction effect was removed improving AIC as shown in this table:

| **call** | **Model** | **df** | **AIC** | **BIC** | **logLik** | **Test** | **L.Ratio** | **p-value** |
|---|---|---|---|---|---|---|---|---|
| With interaction | 1 | 12 | 138.7034 | 166.0234 | -57.35169 | | | |
| Without interaction | 2 | 9 | 136.3427 | 156.8327 | -59.17135 | 1 vs 2 | 3.63932 | 0.3031372 |

#### Model call

```
model <- lme(log2(m_genotype)~Nc+Treatment, data=alldata_m, random=~1|block/plate_ID
,
        weights=varPower(form = ~mut_fitness_genotype * CAA_percent))
```

#### Coefficient table for model 2

| | Value | Std.Error | DF | t-value | p-value |
|---|---|---|---|---|---|
| (Intercept) | 0.391 | 0.2630 | 53 | 1.49 | 0.143 |
| Nc | 0.347 | 0.0549 | 53 | 6.32 | 5.55e-08 |
| TreatmentEF_ATCC194331_none+Controly | 0.667 | 0.1570 | 53 | 4.24 | 8.9e-05 |
| TreatmentMG1655_GFP_Kan1_EF_ATCC194331+Controly | 2.340 | 0.2890 | 53 | 8.09 | 7.93e-11 |
| TreatmentMG1655_GFP_Kan1_none+Controly | 2.500 | 0.3260 | 53 | 7.68 | 3.61e-10 |

#### Coefficient table for model 2

| | Value | Std.Error | DF | t-value | p-value |
|---|---|---|---|---|---|
| (Intercept) | 0.391 | 0.2630 | 53 | 1.49 | 1.43e-01 |
| Nc | 0.347 | 0.0549 | 53 | 6.32 | 1.00e-07 |
| TreatmentEF_ATCC194331_none+Controly | 0.667 | 0.1570 | 53 | 4.24 | 8.90e-05 |
| TreatmentMG1655_GFP_Kan1_EF_ATCC194331+Controly | 2.340 | 0.2890 | 53 | 8.09 | 0.00e+00 |
| TreatmentMG1655_GFP_Kan1_none+Controly | 2.500 | 0.3260 | 53 | 7.68 | 0.00e+00 |

#### ANOVA table for model 2

| | numDF | denDF | F-value | p-value |
|---|---|---|---|---|
| (Intercept) | 1 | 53 | 103.0 | 5e-14 |

| | | | | |
|---|---|---|---|---|
| Nc | 1 | 53 | 1190.0 | <2.2e-16 |
| Treatment | 3 | 53 | 22.4 | 1.65e-09 |

#### Variance and Standard Deviations of model 2

| | **Variance** | **StdDev** |
|---|---|---|
| block = | 0.18100 | 0.4260 |
| plate_ID = | 0.00050 | 0.0224 |
| Residual | 0.00929 | 0.0964 |

#### Weights vary with the following power function

| **mut_fitness_genotype * CAA_percent** |
|---|
| -0.9696005 |

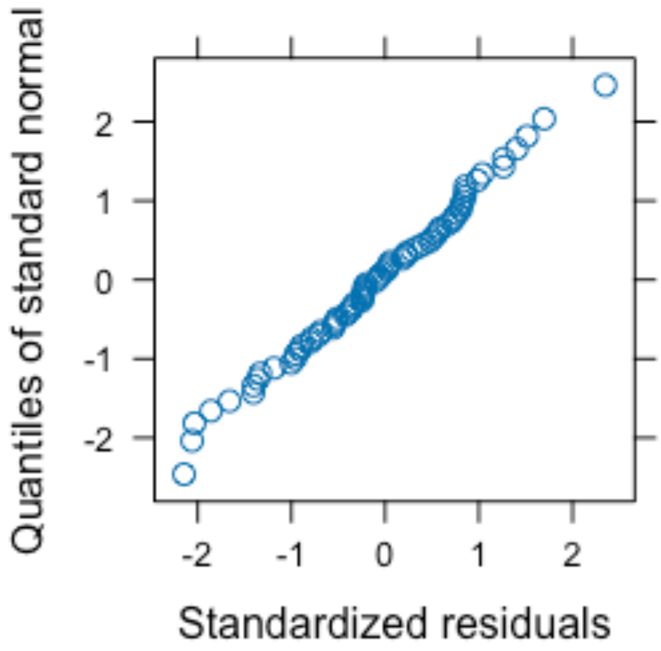


QQ Plot of Deviance Residuals for Model 2

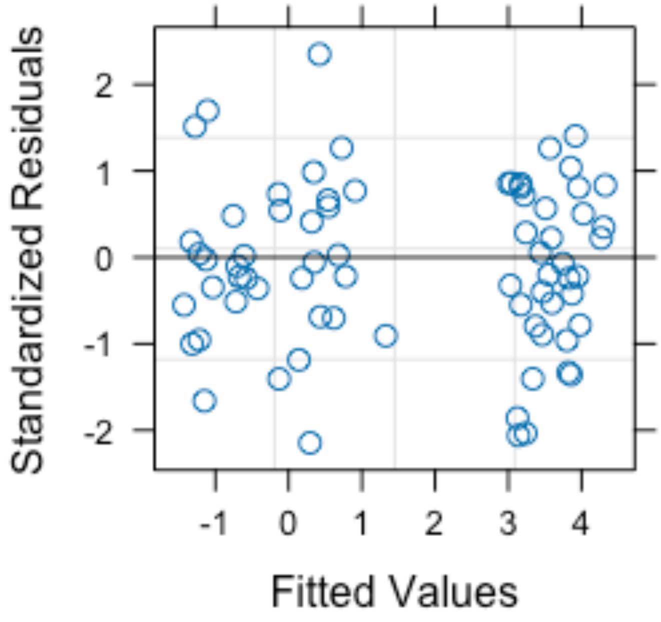


Standardized Residuals vs Fitted Values for Model 2

#### Statistical Model 3

The model shown in Figure 4 (top 4 panels) fits $log_2$ mutation rate as a function of the mean centred $log_2$ final population density of *E. coli* (Dc_ec), *P. aeruginosa* (Dc_pa) and species identity (genotype) allowing for an interaction of species identity with both population density effects. Random effects of individual assay (Pairs) (29 levels) nested within experimental plate (13 levels) nested within experimental block (8 levels), each affecting the intercept, are also fitted.

Predicted values from this model shown as background colours in Figure 4 row 1 and fitted coefficients are shown in Figure 4 row 2.

##### Model call

```
model <- lme(fixed = log2(mutation_rate_genotype) ~ Dc_ec + Dc_pa +
  genotype + Dc_ec:genotype + Dc_pa:genotype, data = PA_coculs,
  random = ~1 | block/plate_ID/Pairs)
```

##### Coefficient table for model 3

| | Value | Std.Error | DF | t-value | p-value |
|---|---|---|---|---|---|
| (Intercept) | 3.4200 | 0.227 | 26 | 15.10 | 2.31e-14 |
| Dc_ec | -0.3710 | 0.170 | 14 | -2.18 | 0.0468 |
| Dc_pa | -0.0734 | 0.262 | 14 | -0.28 | 0.784 |
| genotypePA01 | -0.6620 | 0.294 | 26 | -2.25 | 0.0331 |

| | | | | | |
|---|---|---|---|---|---|
| Dc_ec:genotypePA01 | 0.8570 | 0.236 | 26 | 3.64 | 0.0012 |
| Dc_pa:genotypePA01 | -0.5810 | 0.363 | 26 | -1.60 | 0.121 |

#### ANOVA table for model 3

| | numDF | denDF | F-value | p-value |
|---|---|---|---|---|
| (Intercept) | 1 | 26 | 406.000 | <2.2e-16 |
| Dc_ec | 1 | 14 | 0.948 | 0.347 |
| Dc_pa | 1 | 14 | 3.690 | 0.0753 |
| genotype | 1 | 26 | 0.333 | 0.569 |
| Dc_ec:genotype | 1 | 26 | 18.100 | 0.000238 |
| Dc_pa:genotype | 1 | 26 | 2.570 | 0.121 |

#### Variance and Standard Deviations of model 3

| | Variance | StdDev |
|---|---|---|
| block = | 0.0542 | 2.33e-01 |
| plate_ID = | 0.0000 | 2.65e-05 |
| Pairs = | 0.0000 | 3.98e-05 |
| Residual | 0.9990 | 1.00e+00 |

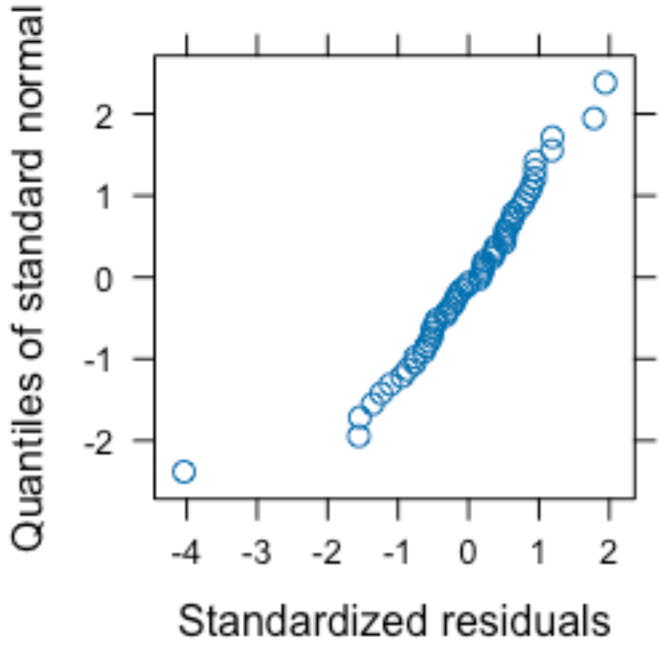


QQ Plot of Deviance Residuals for Model 3

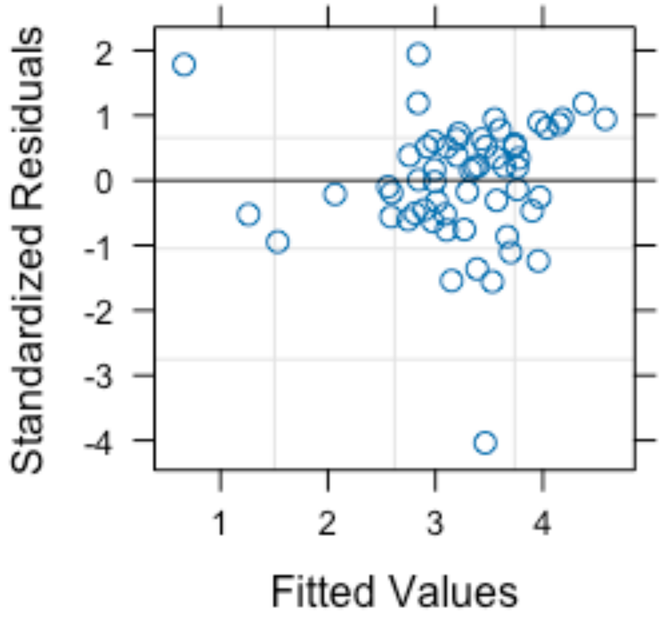


Standardized Residuals vs Fitted Values for Model 3

#### Statistical Model 4

The model shown in Figure 4 (bottom 4 panels) fits $log_2$ mutation rate as a function of the $log_2$ final population size of *E. coli* (NT_ec), *E. faecalis* (NT_ef) and species identity (genotype) allowing for an interaction of species identity with both population size effects. Random effects of individual assay (for which 2 data points, one for each species, are taken) (49 levels) nested within experimental plate (15 levels) nested within experimental block (5 levels), each affecting the intercept, are also fitted.

Predicted values from this model shown as background colours in Figure 4 row 3 and fitted coefficients are shown in Figure 4 row 4.

#### Model call

```
model <- lme(fixed = log2(mutation_rate_genotype) ~ log2(NT_ec) + log2(NT_ef) + genotype + log2(NT_ec):genotype + log2(NT_ef):genotype, data = ModelData, random = ~1 | block/plate_ID/assay_ID)
```

#### Coefficient table for model 4

| | Value | Std.Error | DF | t-value | p-value |
|---|---|---|---|---|---|
| (Intercept) | 11.200 | 11.5000 | 20 | 0.976 | 0.341 |
| log2(NT_ec) | 0.171 | 0.4110 | 8 | 0.415 | 0.689 |
| log2(NT_ef) | -0.463 | 0.0895 | 8 | -5.170 | 0.000856 |
| genotypeMG1655_GFP_Kan | 12.200 | 12.1000 | 20 | 1.010 | 0.325 |
| log2(NT_ec):genotypeMG1655_GFP_Kan | -1.070 | 0.4250 | 20 | -2.510 | 0.021 |
| log2(NT_ef):genotypeMG1655_GFP_Kan | 0.738 | 0.0945 | 20 | 7.810 | 1.68e-07 |

#### ANOVA table for model 4

| | numDF | denDF | F-value | p-value |
|---|---|---|---|---|
| (Intercept) | 1 | 20 | 405.000 | 9.55e-15 |
| log2(NT_ec) | 1 | 8 | 1.840 | 0.212 |
| log2(NT_ef) | 1 | 8 | 0.713 | 0.423 |
| genotype | 1 | 20 | 13.000 | 0.00179 |
| log2(NT_ec):genotype | 1 | 20 | 1.320 | 0.265 |
| log2(NT_ef):genotype | 1 | 20 | 61.000 | 1.68e-07 |

#### Variance and Standard Deviations of model 4

| | Variance | StdDev |
|---|---|---|
| block = | 0.1790 | 0.423 |

| | | |
|---|---|---|
| plate_ID = | 0.0877 | 0.296 |
| assay_ID = | 0.0234 | 0.153 |
| Residual | 0.2670 | 0.516 |

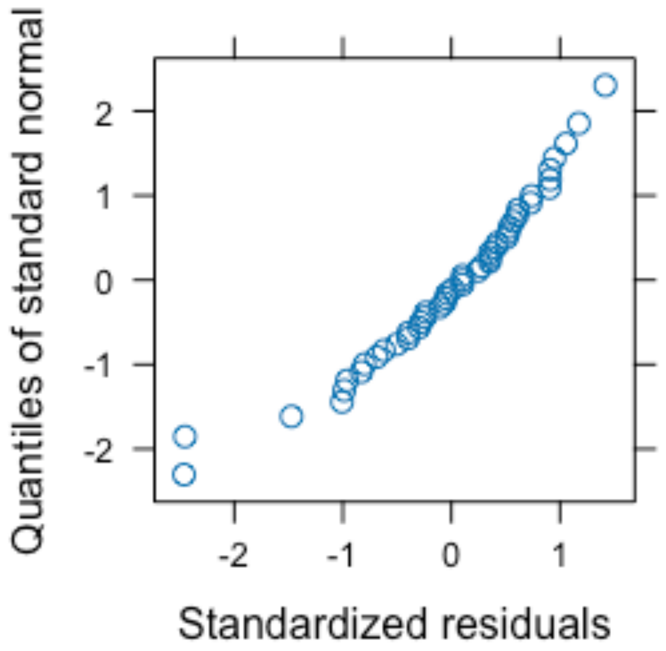


QQ Plot of Deviance Residuals for Model 4

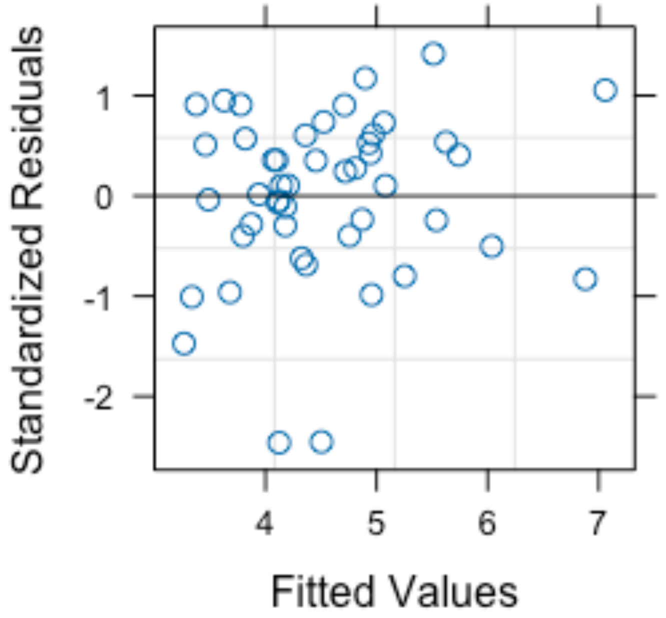

Standardized Residuals vs Fitted Values for Model 4

#### Statistical Model 5

The model shown in Figure S10 fits *E. faecalis* chain length as a function of monoculture vs co-culture growth condition. This count data is fitted by a Poisson distribution using a log link.

##### Model call

```
model <- glm(length ~ condition, Chains, family = poisson(link = "log"))
```

##### Coefficient table for model 5

| | Estimate | Std. Error | z value | Pr(>\|z\|) |
|---|---|---|---|---|
| (Intercept) | 1.580 | 0.0575 | 27.5 | 1.17e-166 |
| conditionmono | -0.235 | 0.0940 | -2.5 | 0.0124 |

##### ANOVA table for model 5

| | Df | Deviance | Resid. Df | Resid. Dev | Pr(>Chi) |
|---|---|---|---|---|---|
| NULL | | | 108 | 175 | |
| condition | 1 | 6.36 | 107 | 168 | 0.0117 |

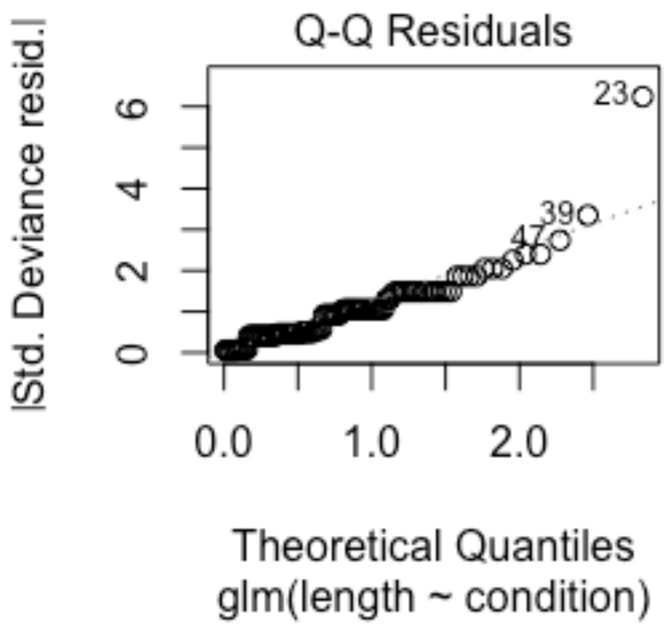


QQ Plot of Deviance Residuals for Model 5

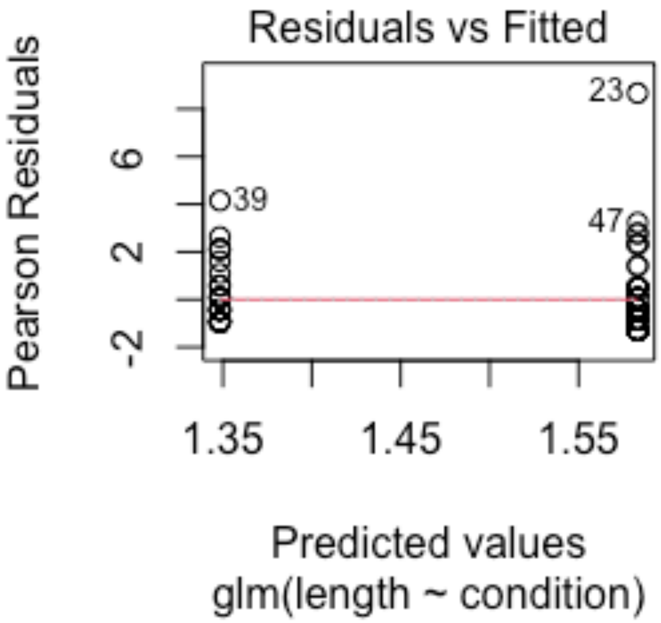


Standardized Residuals vs Fitted Values for Model 5

#### Statistical Model 6

Results presented in Figure 3, Figure 4, Figure S2 and Figure S3 show estimates of the mutation rate to rifampicin resistance (rifR) calculated by coestimation of mutational events per parallel culture and fitness of the wildtype population in reference to the rifR population (‘$\rho$’) (see Methods section ‘Estimation of mutation rates’). In a minority of cases estimation of $\rho$ using the GF method in R package flan fails (2 of 72 in the *Ec* + *Ef* system and 4 of 158 in the *Ec* + *Pa* system). In these cases $\rho$ is estimated based on the successful estimates in other assays as follows:

In the case of *E. faecalis* cultures (in the *Ec* + *Ef* system the are no $\rho$ estimation failures for *E. coli*) the model (model 6.1) takes the mean (on the log scale) of $\rho$ with no fixed effects accounting for random effects of experimental block and plate:

```
model <- lme(log2(mut_fitness_genotype) ~ 1,
    data = EF_GF_data,
    random = ~ 1 | block/plate_ID )
```

##### Coefficient table for model 6.1

| | Value | Std.Error | DF | t-value | p-value |
|---|---|---|---|---|---|
| (Intercept) | 0.0839 | 0.244 | 19 | 0.343 | 0.735 |

##### ANOVA table for model 6.1

| | numDF | denDF | F-value | p-value |
|---|---|---|---|---|
| (Intercept) | 1 | 19 | 0.118 | 0.735 |

#### Variance and Standard Deviations of model 6.1

| | Variance | StdDev |
|---|---|---|
| block = | 0.225 | 4.74e-01 |
| plate_ID = | 0.000 | 5.92e-05 |
| Residual | 0.447 | 6.69e-01 |

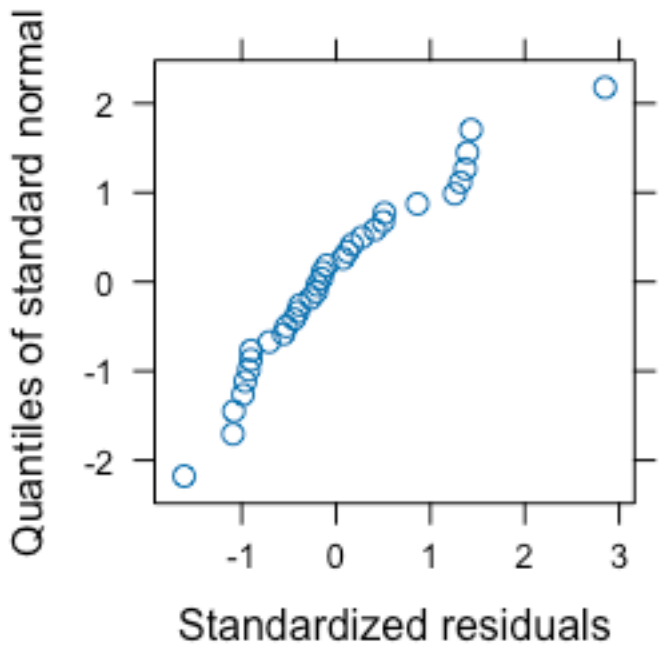


QQ Plot of Deviance Residuals for Model 6.1

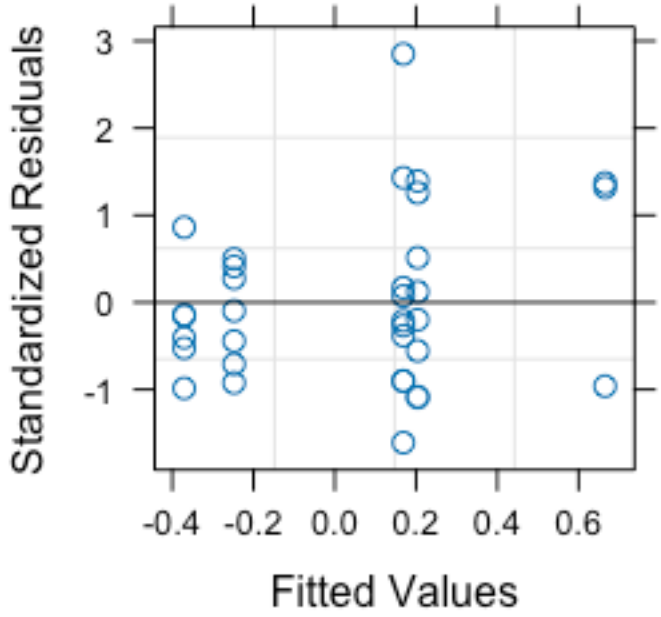


Standardized Residuals vs Fitted Values for Model 6.1

In the case of *E. coli* and *P.aeruginosa* cultures the model (model 6.2) takes the mean (on the log scale) with a fixed effect of treatment (concatenation of focal and co-cultured species) accounting for random effects of experimenter, experimental block and plate:

```
model <- lme(log2(mut_fitness_genotype) ~ treatment,
    data = raw_data2,
    random = ~ 1 | Experimentor/block/plate_ID )
```

#### Coefficient table for model 6.2

| | Value | Std.Error | DF | t-value | p-value |
|---|---|---|---|---|---|
| (Intercept) | 1.400 | 0.391 | 119 | 3.580 | 0.000498 |
| treatmentMG1655_GFP_Kan_PA01 | -0.223 | 0.401 | 119 | -0.556 | 0.579 |
| treatmentPA01_MG1655_GFP_Kan | -1.000 | 0.401 | 119 | -2.500 | 0.0137 |
| treatmentPA01_none | -1.210 | 0.397 | 119 | -3.040 | 0.00291 |
| treatmentPA01_none_LB | -0.881 | 0.480 | 8 | -1.830 | 0.104 |

#### ANOVA table for model 6.2

| | numDF | denDF | F-value | p-value |
|---|---|---|---|---|
| (Intercept) | 1 | 118 | 10.90 | 0.001250 |
| treatment | 4 | 118 | 5.29 | 0.000595 |

#### Variance and Standard Deviations of model 6.2

| | Variance | StdDev |
|---|---|---|
| Experimentor = | 0.0345 | 0.18600 |
| block = | 0.0000 | 0.00014 |
| plate_ID = | 0.0170 | 0.13000 |
| Residual | 1.0100 | 1.00000 |

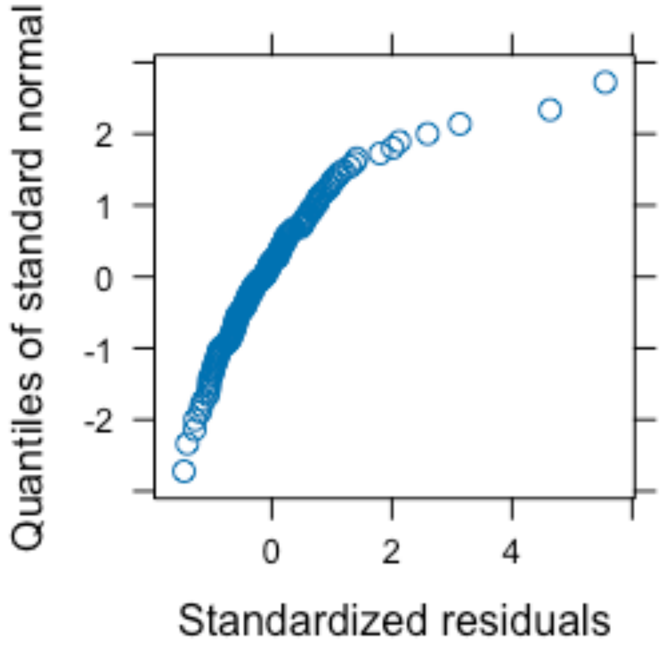


QQ Plot of Deviance Residuals for Model 6.2

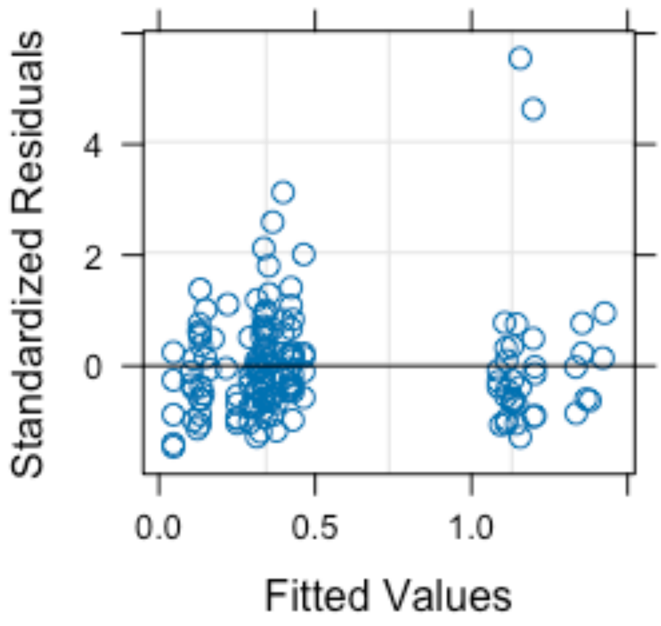


Standardized Residuals vs Fitted Values for Model 6.2
