## Supplementary figures and images for "Cohabiting microbes can impact one another’s mutation rates creating a mutational eco-evo feedback loop"

### Graphical Abstract

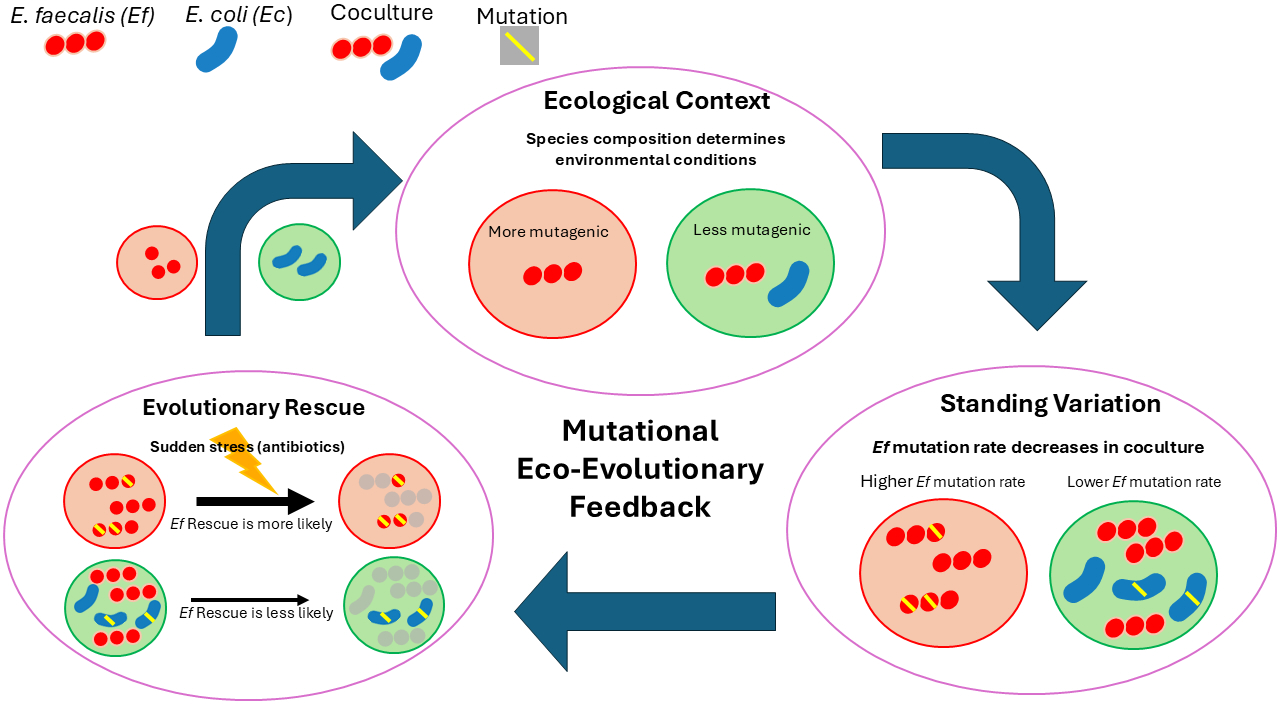
